# CORGIAS: identifying correlated gene pairs by considering evolutionary history in a large-scale prokaryotic genome dataset

**DOI:** 10.1101/2025.05.07.652372

**Authors:** Yuki Nishimura, Kimiho Omae, Kento Tominaga, Wataru Iwasaki

## Abstract

The recent expansion of prokaryotic genomes reveals many ortholog groups (OGs) whose function cannot be inferred from conventional, sequence similarity-based annotation methods, especially in metagenome-assembled genomes. Phylogenetic profiling is one of the promising methods to annotate these OGs, by identifying functional relationships of OGs using co- or anti-occurrence of OGs distributions, not sequence similarity. Here, we proposed two new phylogenetic methods for large-scale data, Ancestral State Adjustment (ASA) and Simultaneous EVolution test (SEV), which consider the ancestral state of gene presence/absence. In evaluations using three distinct prokaryotic datasets, ASA and SEV showed better or comparable performance to both established and recently proposed methods for large-scale data. We compared the functionally related genes detected by each method and found that SEV and its predecessor can identify slowly evolving genes, such as housekeeping genes. In contrast, ASA and its predecessors can detect functionally related genes that tend to be gained or lost in a fixed-order, indicating a strong evolutionary constraint that provides clues for functional prediction. Using matrix multiplication, we showed that SEV is scalable in the latest genome databases.

## 1. Introduction

Advancements in sequencing technology have allowed access to a number of genomes of uncultured prokaryotes, including those forming large lineages that occupy the unignorable parts of prokaryotic diversity (1, 2). These ‘microbial dark matter’ harbor many ortholog groups (OGs) showing no similarity in amino acid sequences to those functionally annotated in previous studies (3–6). As functionally unknown OGs in microbial dark matter are estimated to constitute up to 50% of their genomes (7), it is important to reveal their function to understand their ecology and evolution and to apply genetic resources to industry, agriculture, and medicine.

Phylogenetic profiling is a method to find the co-evolved OGs that are expected to be functionally related from a matrix indicating the presence/absence or copy number of genes in OGs, so-called profile, in many genomes (8). Once a set of correlated OGs has been identified, they are expected to be functionally related. Thus, the function of unannotated OGs can be inferred from the annotations of other OGs in the correlated gene set (9–14).

In addition to positively correlated OGs, negatively correlated OGs can help understand the function of OGs (15–19). Recently, we proposed an approach that uses negatively correlated OG pairs for functional prediction as ‘contrapositive genetics’. As a proof of concept, we showed that the function of unannotated OGs frequently found in the genome lacking those for replicative helicase loaders work as helicase loaders (20).

One of the challenges in phylogenetic profiling is distinguishing co- and anti-occurring OGs that are functionally related from those just inherited from a common ancestor and have no functional relation. For example, the presence/absence of two pairs of OGs is shown in Fig. 1A. In both cases, the two OGs co-occur. However, in the upper pair, the co-occurrence emerged in the multiple independent lineages, whereas in the lower pair, it emerged only once in the common ancestor. Therefore, the upper pair appears to have co-evolved while the lower does not. Nevertheless, when simply counting the number of genomes by presence/absence state of pair OGs (the naive method), two pairs cannot be statistically distinguished as they yield the same contingency table (Fig. 1A). Several methods have been developed to overcome this problem. These methods can be classified into two types: model-based and heuristic. A model-based approach explains the OG distribution patterns based on a statistical evolutionary model. For example, in Pagel’s method (21), likelihoods of the paired OG distribution in the phylogenetic tree under dependent (co-evolved) and independent models are evaluated. Then, a likelihood ratio test is performed to assess whether the paired OG is co-evolved or not. Although Pagel’s method and other model-based approaches are designed to evaluate paired OG (22, 23), the evolCCM can model interactions including more than three OGs (24). However, model-based approaches are computationally expensive and difficult to apply to large prokaryotic datasets that can include thousands or tens of thousands of species and OGs these days (25–27).

**Figure 1.**
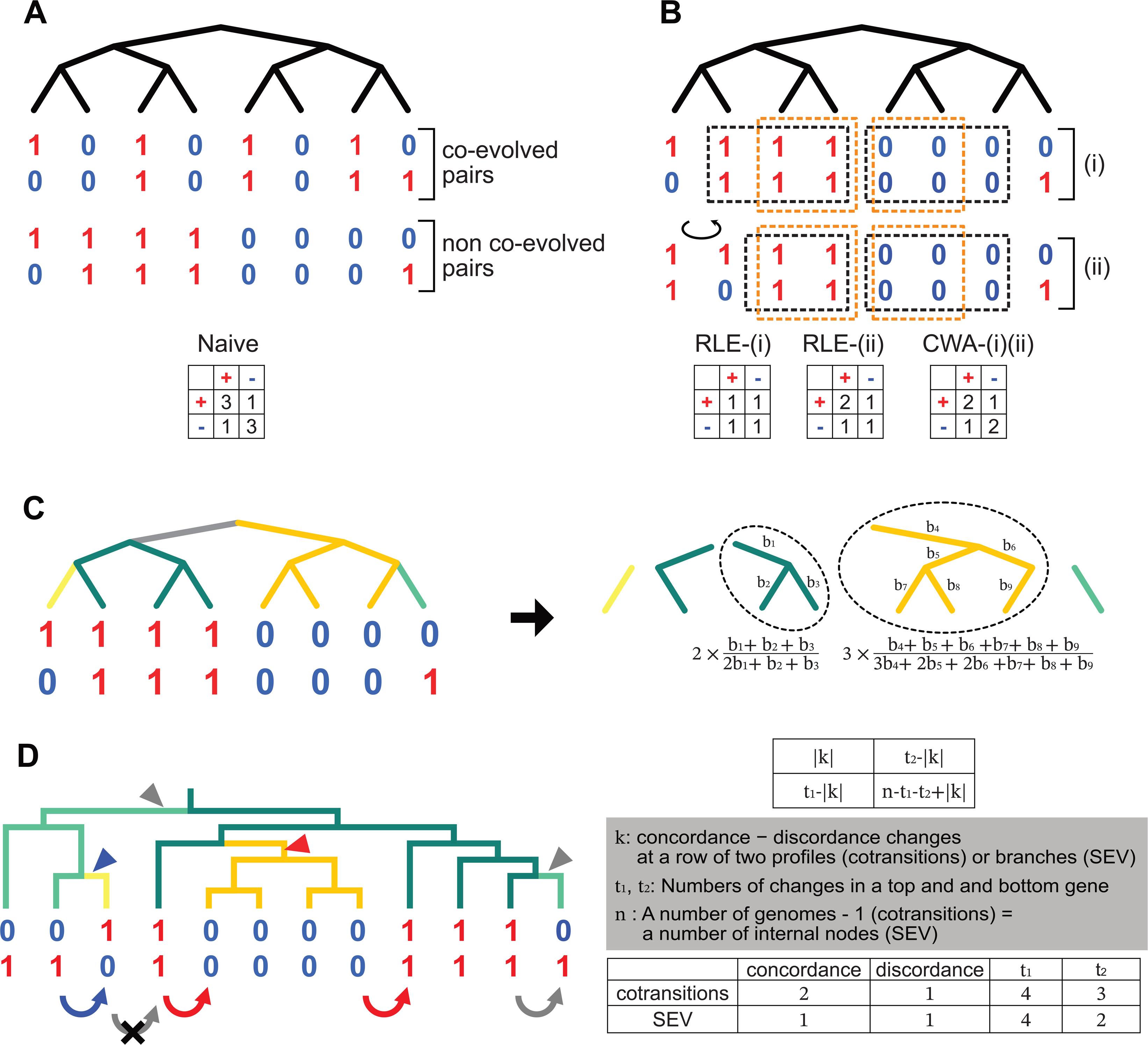
Schematic diagram of phylogenetic profiling. (A) Phylogenetic profiles of a non-co-evolved (top) and a co-evolved gene pair (below). Whether a pair of genes is co-evolved or not is determined by the statistical significance of the co-occurrence. In the naive phylogenetic profiling methods, both co-evolved and non-co-evolved pairs are tested with the same contingency table (bottom). Thus, they are statistically indistinguishable. (B) Application examples of RLE and CWA to the non-co-evolved pair in (A). Profiles surrounded by black and orange dotted squares are compressed to one profile in RLE and CWA, respectively. In the middle, the leftmost two branches and their profiles of the top figure are rotated. The resultant contingency tables are shown at the bottom. (C) The schematic diagram of ASA. A phylogenetic tree is divided into subtrees based on the reconstructed ancestral states of a paired-gene profile (left panel). The tree branches are colored by the result of ancestral state reconstruction. A number of genomes in a subtree with more than two leaves (surrounded by a dotted circle) are submitted to correction, according to evolutionary time (right panel). The corrected numbers of genomes are shown below the dotted circles, where b_n_ represents the branch length of node n. In the method proposed by Ruano-Rubio et al., each subtree is counted as one. (D) The schematic diagram of cotransitions and SEV. The curved arrows indicate presence/absence changes counted in cotransitions, while triangles indicate those counted in SEV. Concordance and discordance changes are shown in red and blue, respectively, while gray represents changes only in either of the genes. In cotransitions, consecutive changes are ignored as indicated by the gray arrow with a cross. The significance of the correlation of a pair of genes is tested by using the contingency table in the top right panel. The notation in the contingency table is shown in the middle right panel (Although Dembech et al. designated n as the number of genomes, the number of genomes minus one is correct because transitions can be placed between genomes). The number of concordances, disconcordances changes, t_1_, t_2_ in cotransitions, and SEV are shown in the bottom right panel. Note that the concordance gene gain occurred only once (red triangle) but was counted as two transitions in cotransitions.

In contrast, heuristic methods compare profiles of paired OGs, trying to reduce the phylogenetic effect. For example, in Run-length encoding (RLE, Fig. 1B), a pair of profiles is firstly ordered along genomes in a species tree, and consecutive chunks of profiles sharing the presence/absence state (“runs”) are compressed and counted as one (28). Although RLE is so fast as to be used to find co-evolved pairs in approximately thirty thousand bacterial genomes (29), it has two problems: Branch rotation around internal nodes does not change the tree topology, but RLE makes different results from the same data (Fig. 1B), and genomes in the same run are not necessarily evolutionarily close. These problems can be avoided by clade-wise adjustment (CWA; Fig. 1B), which compresses the state-sharing profiles only when they form a monophyly (30). Still, it cannot compress the profiles sharing the states derived from a common ancestor and form a paraphyletic or polyphyletic clade. Ruano-Rubio et al. utilized ancestral state reconstruction to select genomes that are compressed more rationally (31). However, these ‘weighted methods’ ignore the evolutionary time the genomes have undergone as they compress the profiles sharing presence/absence state into just one. A corrected number of profiles should be determined considering the evolutionary time.

In contrast to weighted methods, Dembech et al. proposed a novel heuristic approach, cotransitions, in which profiles are firstly ordered along a phylogenetic tree as well as RLE but focus on the number of presence/absence changes in rows of profiles (12). Then, a statistical test using the numbers of genomes and changes is conducted to detect a correlation in the paired OG (Fig. 1D). Although cotransitions is highly scalable, the results can be changed by branch rotation as well as RLE. In fact, the authors analyzed the same dataset in three different orientations and reported that only 22% of the detected co-evolved pairs overlapped among the analyses (12). In addition, gain/loss events cannot be counted correctly. For example, in Fig. 1D, the simultaneous gain of paired OGs occurs as indicated by a red triangle, but this is counted as two as indicated by red two arrows (Fig. 1D). Note that not all gain/loss events are duplicated because the consecutive changes are ignored (the gray arrow with a cross in Fig. 1D).

Here, we propose two new heuristic approaches that utilize the reconstructed ancestral states of the profile to evaluate the evolution of OGs properly and obtain robust results against branch rotation. One is the Ancestral State Adjustment (ASA), a variant of the weighted methods that calculates the corrected number of genomes by considering ancestral states and evolutionary time. The second is a Simultaneous EVolution test (SEV) modified from cotransitions. The two proposed methods showed better or comparable performances to the previous heuristic methods and those implemented in a recent tool, EvoWeaver (32), in evaluation using three distinct prokaryotic datasets. SEV also showed high scalability that can be applied to large-scale datasets. Comparison of the co-evolved OG pairs detected by each method revealed that SEV and cotransitions can find pairs that have been gained/lost infrequently and those that are unevenly distributed among the genomes. In contrast, pairs detected only by weighted methods tend to have a specific order of gene gain/loss. These characteristics can be useful for estimating the functions of co-evolved OGs identified by phylogenetic profiling. ASA and SEV are freely available in a Python package called CORelated Genes Identifier by Considering Ancestral State (CORGIAS), with the reimplementation of RLE, CWA, and cotransitions at https://github.com/ynishimuraLv/corgias.git.

## 2. Materials and Methods

### 2.1 Ancestral state adjustment (ASA)

We introduced an ASA that corrects the number of paired OG profiles sharing presence/absence states from a common ancestor, considering the evolutionary time. The ancestral states of the profiles at each node in a species tree should be given, but ambiguous states are allowed. The methods inferring the ancestral state are arbitrary; however, in this study, we adopted a maximum-likelihood method using pastML (33). In the first step of ASA, a tree is divided into subtrees based on the estimated ancestral states of the paired OGs (Fig. 1C, left panel). Each subtree has the same profile of a paired OG from the root to the leaves. This means that all descendants inherited the profile that emerged at the root of the subtree, and each subtree is subject to adjustment, corresponding to ‘run’ in RLE. The root of the subtree is identified by tracing back from the leaves (extant species) to their ancestral nodes until the ancestral profile is changed or uncertain. Then, the corrected number of profiles, M_c_, in the respective subtrees is calculated by the following equation:

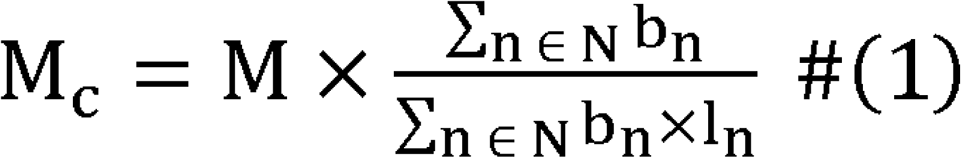

where M is the number of genomes in the subtree; N is the set of all leaves and internal nodes in the subtree; b_n_ is the branch length between node n and its ancestral node; and l_n_ is the number of leaves derived from node n, respectively. The second term (fraction) is equivalent to the sum of the branch lengths in the subtree divided by the sum of the branch lengths in the star tree, converted from the subtree. The denominator is assumed to represent the evolutionary time if they have evolved independently. That means that the number of profiles is adjusted by the ratio of dependent evolutionary time to that of independent. Formula (1) implies that the corrected number of profiles decreases as the number of leaves increases, and the internal branch lengths representing the dependent evolutionary time are longer. The corrected numbers of profiles are summed by the presence/absence state of the paired OG and subjected to Fisher’s exact test to determine the significance of their correlation. When the sum of the corrected number of genomes is not an integer, it is rounded up before performing Fisher’s exact test.

### 2.2 Simultaneous EVolution test (SEV)

SEV is very similar to cotransitions (12) but focuses on presence/absence changes at the branch, not in rows of profiles, as is counted in cotransitions (Fig. 1D). A concordance change, in which both OGs are simultaneously gained or lost between the parent and its child nodes, is a sign of co-evolution. In contrast, a discordance change in which one OG is gained while the other is lost at the same branch indicates contrapositive evolution, in which two OGs have evolved to avoid each other. SEV is also similar to the “Simultaneous Score” in treeWAS (34), but the statistical test was inspired by cotransitions. As with ASA, SEV requires a phylogenetic tree and reconstructed ancestral presence/absence states of the profiles. However, the parallel gain or loss of an OG at the two branches derived from their common parent node is prohibited because of the constraint in the statistical test (see below). The easiest method satisfying this constraint is the maximum parsimony method. A one-tailed Fisher’s exact test from a 2 × 2 contingency table below can determine whether an OG pair is correlated.

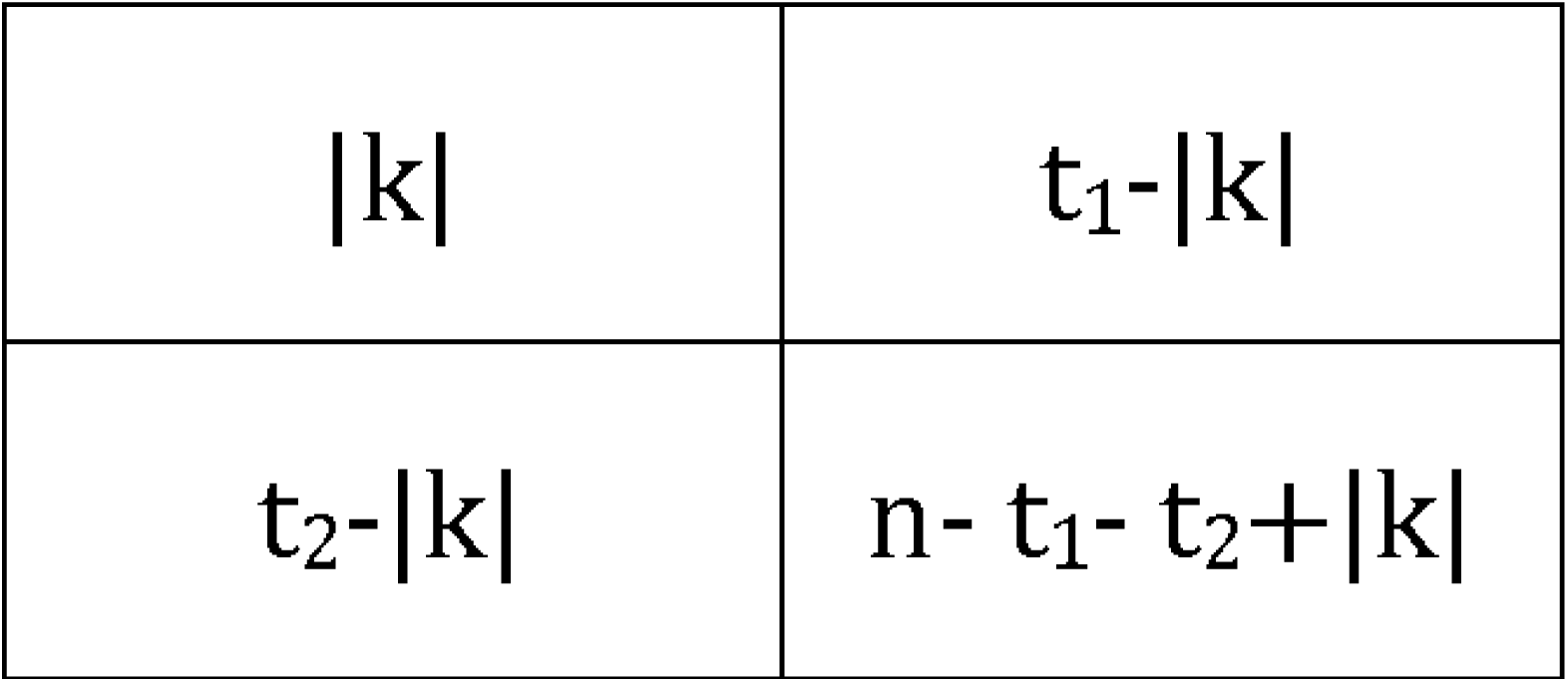

where k is the difference in the number of concordance and discordance changes, t_1_ and t_2_are the numbers of presence/absence changes in each OG, and n is the number of internal nodes of the tree. Under the constraint mentioned above, n is not the number of branches, because the parallel OG gain/loss at a node is never inferred by the maximum parsimony method. When the ancestral state is uncertain at the nodes, the change is regarded as not having occurred at the branches extending from the nodes. When the obtained p-value is significant and k is positive, the pair OG can be regarded as co-evolved, whereas a negative k indicates contrapositive evolution. k for all the paired OG combinations in the dataset can be calculated by multiplying a matrix comprising the vector representing each OG gain/loss at the branch, thus providing scalability to the SEV. In the matrix, 1 and −1 indicate the gain and loss of an OG, respectively, while 0 indicates no change. When the ancestral state of the node is uncertain, 0 is placed at the corresponding position in the vector. Notably, although k in cotransitions can, in principle, be calculated using matrix manipulation, the authors’ implementation does not employ such an approach. Thus, we reimplemented the cotransitions to utilize matrix manipulation for performance and runtime evaluation. The validation of reimplementation is available in https://github.com/ynishimuraLv/corgias_data.

### 2.3 Preparing the prokaryotic datasets for performance evaluation

The proteome data inferred from representative genomes belonging to three distinct lineages, Order Pseudomonadales (gram-negative), Order Mycobacteriales (gram-positive), and Domain Archaea, were downloaded from GTDB Release 214 (35). The qualities of the downloaded proteomes were assessed using checkM2 (36). Proteomes with less than 90% completeness or > 10% contamination were excluded from the analysis. The remaining 2, 181, 1, 627, and 1, 922 proteomes of Pseudomonadales, Mycobacteriales, and Archaea, respectively, were used for further analysis.

All proteins in the datasets were submitted to COGclassifier (https://github.com/moshi4/COGclassifier), which assigns proteins to Clusters of Orthologous Genes (COGs (37)). The results were used to prepare profiles for each dataset; A COG is regarded as ‘present’ in a genome if at least one gene in the genome was assigned to that COG. Note that not all genes in the genomes were assigned to COG. As it is difficult to identify co-evolved OGs contained in almost all or few genomes by phylogenetic profiling, only COGs found in 1-99% of the genomes were retained, resulting in 3, 232, 2, 814, and 3, 307 COGs in Pseudomonadales, Mycobacteriales, and Archaea, respectively. A phylogenetic tree of archaea with 1, 922 species was prepared from the ar53_r214.tree provided by GTDB by subtracting the genomes of the discarded proteomes. Phylogenetic trees of Pseudomonadales and Mycobacteriales were prepared as well, but from the bac120_r214.tree. Ancestral state reconstructions were performed on each COG profile and tree using pastML (33) by MPPA (the maximum likelihood method recommended by the developers) for ASA and ACCTRAN for SEV, because they always showed the best area under the precision-recall curve (PRAUC) in all datasets (Supplementary Table S1).

To compare the performance of each method, the grand truth set consisting of truly functionally related COG pairs is required. We used the COG pair scores derived from the STRING database v12 (38) as in the previous study (16). As STRING scores were calculated from gene neighborhood conservation, gene fusion, co-expression, protein interaction experiments, other databases, text mining, and occurrence patterns, the scores were recalculated without occurrence to make the score independent of co-occurrence. COG pairs with a recalculated score larger than 0.9 were regarded as positive, as it is the lower limit of the highest confidence of functional association in the STRING database; all others were considered negative. The number of COG pairs scored in STRING and the positive pairs in the datasets are summarized in Table 1. It should be noted that the majority of COG pairs (∼80%) do not have the STRING scores as shown in Table 1 and these COG pairs were ignored from the performance evaluation. The distributions of the STRING score used in this study were shown in Supplementary Figure S1. We also justified thresholds of 0.9 for functional correlation by confirming that the union of the top-ranking pairs across all examined methods before the True Positive Rate (TPR) fell below 50% showed score distributions biased toward values above 0.9 (Supplementary Figure S1).

**Table 1.** The summary of prokaryotic datasets analyzed in this study. The ratio of scored COG pairs to those of all pairs are shown in parentheses.

| Dataset | Genomes | COGs | Scored pairs | Positive pairs |
| --- | --- | --- | --- | --- |
| **Pseudomonadales** | 2,180 | 3,233 | 1,069,619 (20.5%) | 5,870 |
| **Mycobacteriales** | 1,675 | 2,814 | 846,124 (21.4%) | 4,040 |
| **Archaea** | 1,920 | 3,307 | 1,297,337 (23.7%) | 6,003 |

### 2.4 Preparing the simulation datasets for evaluation of runtime performance

Two groups of datasets were simulated to compare the runtimes of each method. One group contained six datasets with 1, 000 genomes and varying numbers of OGs. To this end, a phylogenetic tree was generated with the “pbtree” function in phytools (39) and used for simulating 1, 000, 2, 000, 4, 000, 8, 000, 16, 000, and 32, 000 OG pairs by evolCCM (24). The evolCCM uses five parameters to generate pairs of OGs. One of these parameters is the coefficient of interaction between the two OGs. To generate co-evolved OG pairs, the coefficients were sampled from a uniform distribution ranging from 0.2 to 0.75 while setting the coefficient to zero for non-co-evolved OGs. Co-evolved and non-co-evolved pairs were generated at a ratio of 1:99. Two parameters representing the intrinsic rate of each OG were randomly sampled from a uniform distribution from −0.5 to 1. The last two parameters indicated half the difference between the gain and loss rates of each OG and were sampled from a uniform distribution from −0.5 to 0.3. Profiles with 2, 000-64, 000 OGs were prepared by concatenating the simulated OG pairs within the datasets. The second group contains six datasets with 4, 000 OGs and various numbers of genomes (500, 1, 000, 2, 000, 4, 000, 8, 000, and 16, 000). Phylogenetic trees and profiles were prepared in the same way as in the first group, but with different numbers of OG pairs and genomes.

The runtimes for calculating the number of genomes (in the naive method), the corrected number of genomes (in RLE, CWA, and ASA), or k, t1, t2, and n (in cotransitions and SEV) were measured from a phylogenetic tree, a profile, and the inferred ancestral state of OGs in each dataset of the two groups. As the naive method, cotransitions, and SEV included multiplications of matrices, calculations were performed both with and without a GPU. All runtime measurements were executed on a computer with 56 CPUs of 2.60 GHz, 512 GB of RAM, and a single NVIDIA RTX A6000. Analyses that lasted longer than one day were terminated.

### 2.5 Performance comparison with EvoWeaver

We compared our proposed methods (ASA and SEV) with the four phylogenetic profiling methods implemented in a new tool, EvoWeaver (32). For performance comparison, the three prokaryotic datasets were analyzed by the phylogenetic profiling methods in EvoWeaver. For runtime comparison, we focus on one of the methods, G/L Distance because it showed the most promising performance in our dataset. As EvoWeaver does not perform phylogenetic profiling parallelly, we did it with the ‘parallel’ package (https://dept.stat.lsa.umich.edu/~jerrick/courses/stat701/notes/parallel.html). The runtimes for the dataset with more than 32, 000 OGs cannot be measured due to the shortage of RAM.

## 3. Results

### 3.1 Performances in the Prokaryotic Datasets

Three distinct prokaryotic datasets containing 1, 600-2, 200 genomes were prepared: Order Pseudomonadales (gram-negative bacteria), Order Mycobacteriales (gram-positive bacteria), and Domain Archaea. All combinations of COG pairs in the datasets were evaluated using naive, RLE, CWA, ASA, cotransitions, and SEV. As the datasets are unbalanced, with a few positive (co-evolved) pairs compared to negative (non-co-evolved) pairs (Supplementary Table S2), the performances were evaluated based on the PRAUC. We also compared the number of detected positive pairs at the true positive rate (TPR) varying from 0.9 to 0.5.

The cotransitions and SEV (hereafter, transition methods) showed the best PRAUC in all three datasets (Fig. 2A). The number of detected positive pairs of either of transition methods were also best in all datasets at all TPRs (Fig. 2B) and covered more than 80% of the pairs detected by all methods, except the Mycobacteriales datasets at TPR=0.9 (Supplementary Tables S2-S4). As the results under this condition do not reflect the general trends, we excluded them from the following comparison. Among transition methods, SEV showed greater than or competitive performance to cotransitions in terms of PRAUC and the number of detected positive pairs (Fig. 2B and Supplementary Table S2-S4). In addition, the number of positive pairs detected only by SEV was several times larger than those detected only by cotransitions in most cases, although not always (Supplementary Figure S2). Considering the above performance and robustness against branch rotation, SEV can be regarded as an improved version of cotransitions.

**Figure 2.**
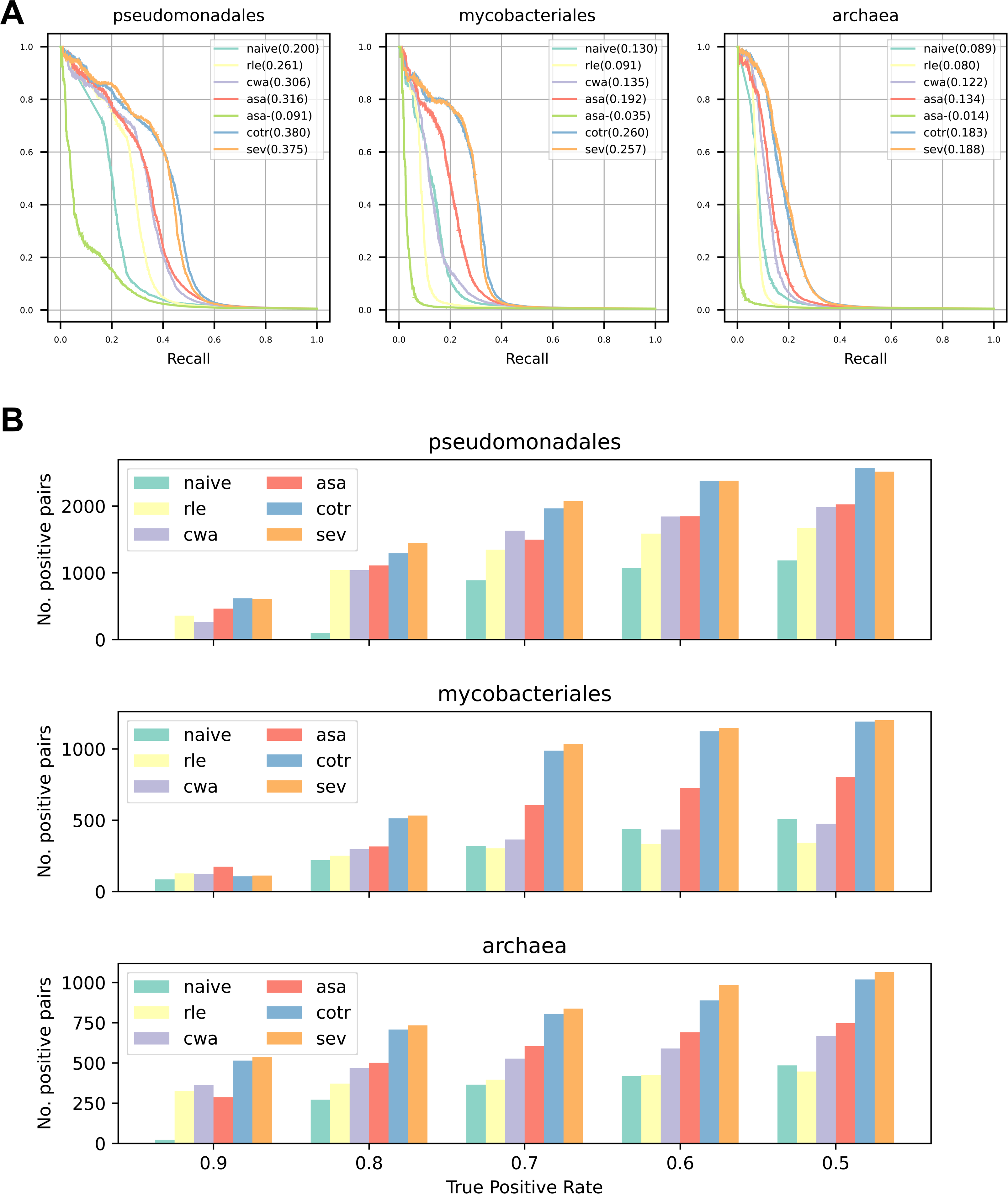
Performance comparison of phylogenetic profiling methods using prokaryotic datasets. (A) Precision-Recall (PR) curves of each method. Areas under the PR curve (PRAUCs) are shown in parentheses. ‘asa-’ indicates ASA without branch length adjustment. (B) The number of detected positive pairs of each method at various true positive (TPR) thresholds.

We postulated that the advantage of SEV is derived from the precise count of gain/loss as the number of gain/loss is duplicated in cotransitions. To inspect this hypothesis, we compared the evolCCM (24) coefficient for the positive detected pairs by SEV but not by cotransitions, and vice versa. The coefficient of evolCCM determines the transition rate between the presence/absence states of the paired OG, the larger the coefficient, the faster the transition rate from the current state to the other states. As a result, the coefficients of the pairs detected by SEV but not contransitions tend to be enriched lower than the opponents (Fig. 3A and Supplementary Figure S3). These results indicate that the precise counts of gain/loss events considering ancestral states contribute to the detection of slowly evolving OGs.

**Figure 3.**
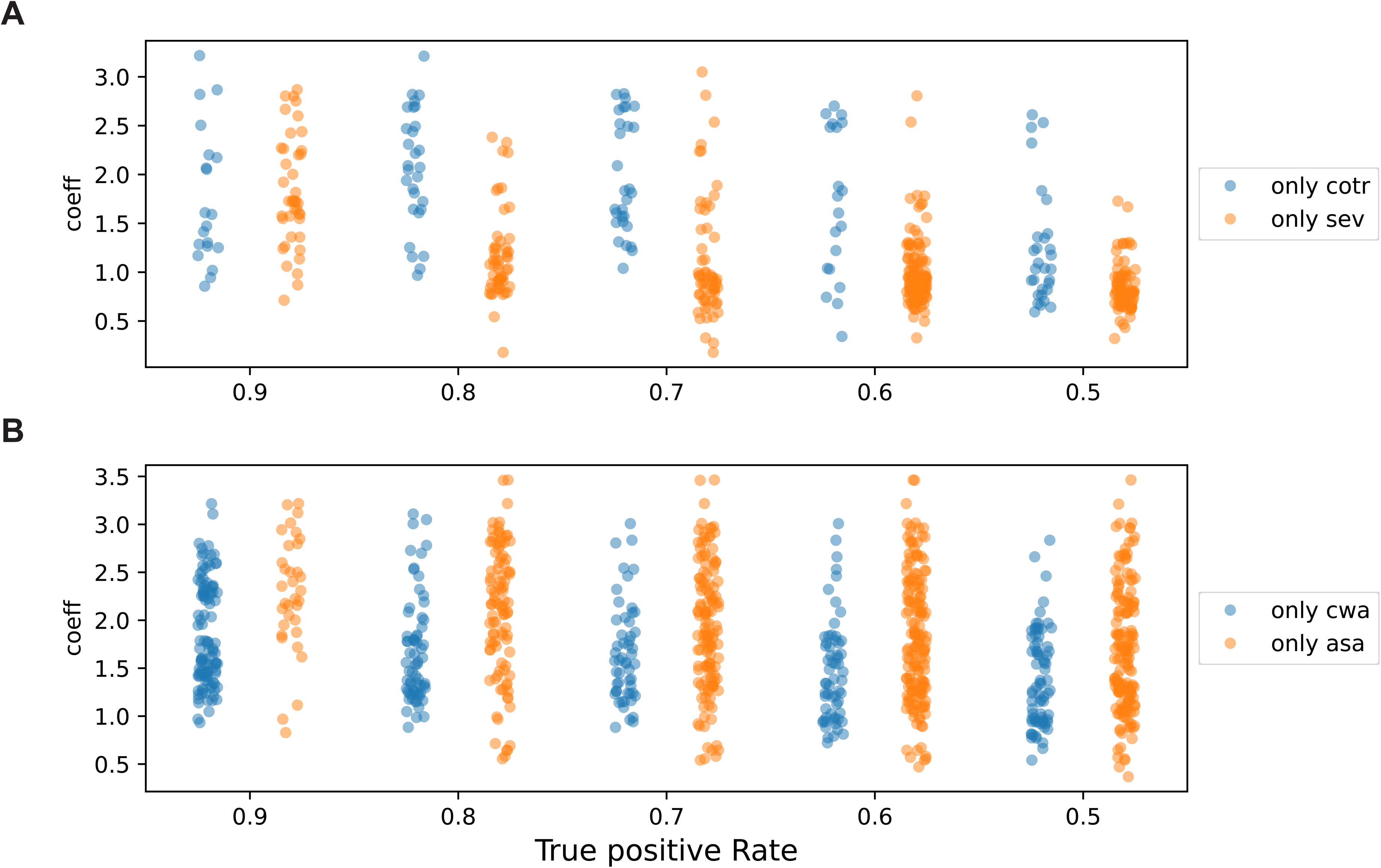
Comparison of the proposed phylogenetic profiling methods in this study and those in the previous studies. (A) The evolCCM coefficient distribution of pairs detected by cotransitions (cotr) but not SEV, and vice versa, in the Archaea dataset at various true positive ratios (TP). (B) Same as (A), but for pairs detected by CWA but not ASA, and vice versa. Comparison in the Pseudomonadales and Archaea datasets are shown in Supplementary Figures S3 and S4.

Among the weighted methods, ASA always showed the best PRAUC (Fig. 2A). It is worth noting that correcting the number of profiles by considering branch lengths is critically important in ASA because when each subtree is counted as one, the PRAUC drastically decreases (Fig. 2A). Although the differences from the second-best method (CWA) were small in the Pseudomonadales and Archaea datasets, ASA often detected more positive pairs than any other weighted method in all datasets (Fig. 2B and Supplementary Tables S2-S4). In addition, CWA results were unstable among the datasets. For example, the ratio of pairs detected by CWA to those detected by either method in the Mycobacteriales dataset was less than half of that in the Pseudomonadale dataset. Similar to CWA, RLE decreased the number of detected positive pairs and was inferior even to the naive method in the Mycobacteriales dataset (Fig. 2B and Supplementary Table S4), suggesting that the performance of CWA and RLE depend heavily on the datasets. In contrast, ASA showed stable performance in all three datasets (Supplementary Table S2-S4), demonstrating the importance of considering the ancestral state and branch lengths. Considering these results, ASA can be regarded as an improved version of the previously proposed weighted methods.

CWA requires monophyly to compress profiles but ASA does not, implying that the pairs detected only by CWA are more conserved and not frequently gained/lost than those detected only by ASA. Indeed, the pairs detected by CWA but not ASA tend to have lower evolCCM coefficients than vice versa (Fig. 3B and Supplementary Figure S4). The constraint of RLE is weaker than that of CWA but the difference in coefficient distribution from ASA is similar to or biased lower than CWA, probably due to RLE’s low precision Supplementary Figure S5).

### 3.2 Transition methods can detect slowly evolving and unevenly distributed OG pairs

The PRAUCs of all weighted methods and the number of detected positive pairs were far from those of the transition methods in all datasets (Fig. 2 and Supplementary Tables S2-S4). Even if the positive pairs detected by the weighted methods accumulated, each transition method still detected more positive pairs (Supplementary Tables S2-S4). The number of pairs detected by either of the weighted methods, but not by either of the transition methods (i.e., pairs only detected by the weighted methods), was less than 10% of all detected pairs (Supplementary Tables S5-S7). In contrast, 20-40% of the pairs detected by either of the transition methods were not found by any weighted method (Supplementary Table S5-S7). To clarify the characteristics and advantages of transition methods over weighted methods, we again compared evolCCM coefficients between all positive pairs detected by either the weighted or transition methods.

We first analyzed the coefficient distribution at TPR=0.7. Comparing the coefficient distribution between the pairs detected only by the weighted and transition methods, those of the transition methods had peaks at low values (1-1.5) in all datasets (Fig. 4A), suggesting that the transition methods can detect slowly evolving OG pairs. In other words, transition methods can detect more conserved OG pairs than weighted methods. For example, many pairs of COGs encoding ribosomal proteins and translation factors (classified as COG category J) were detected only by the transition methods and had coefficients of approximately 1 in all three datasets (Supplementary Fig. S7). As ribosomes are indispensable for cellular life (40), it is reasonable that their gains and losses rarely occur, resulting in low coefficients. Owing to the high conservation of ribosomal proteins, almost all genomes have COGs detected by the transition methods; the ribosomal protein COGs consisting of the positive pairs only detected by transition methods are possessed by 96, 84, and 91% of Pseudomonadales, Mycobacteriales, and Archaea genomes in the dataset, respectively. These high possession rates lower the corrected number of profiles, rendering them undetectable by weighted methods.

**Figure 4.**
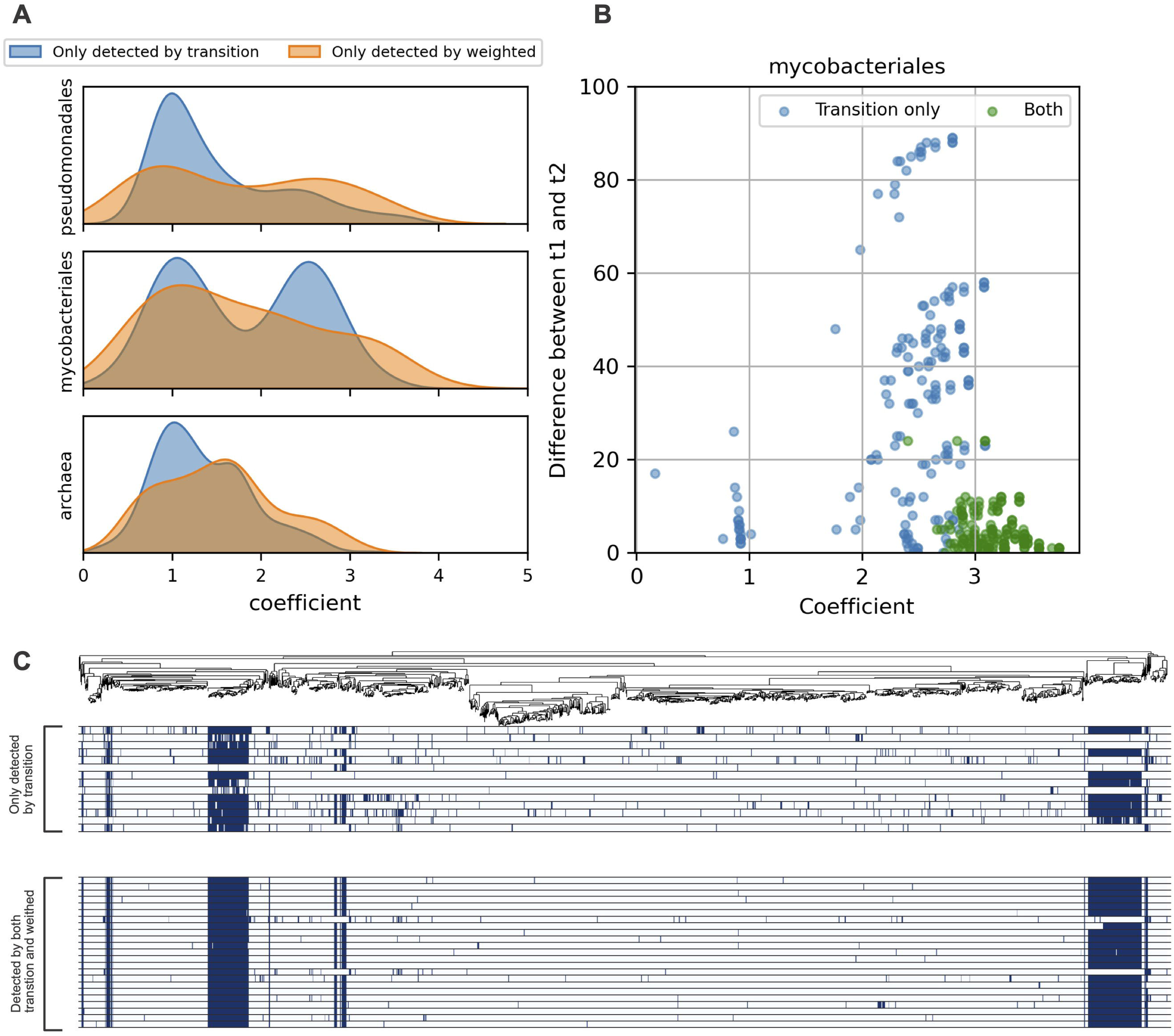
Characteristic of transition methods. (A) The evolCCM coefficient distribution of the co-evolved pairs only detected by transition methods (blue) and weighted methods (orange) at a True Positive rate (TPR) = 0.7. (B) The scatter plot of the coefficient (x-axis) and the differences between t_1_and t_2_ in co-evolved pairs of COGs belonging to category N (y-axis) in the Mycobacteriales datasets at a TPR = 0.7. t_1_ and t_2_ are the number of presence/absence state changes in each COG of the pairs across the phylogenetic tree. Blue dots indicate the co-evolved pairs only detected by transition methods, while green ones indicate pairs detected by both transition and weighted methods. Note that no pairs were detected only by the weighted methods, except for two with large differences between t_1_ and t_2_. The plots for the other categories in all datasets are shown in Supplementary Figs. S7-S9. (C) Phylogenetic distribution of co-evolved COGs belonging to COG category N in Mycobacteriales at TPR = 0.7. The top panel shows the species tree of Mycobacteriales used in this study. The middle and bottom heatmap shows the presence/absence of the genes only detected by transition methods and those detected by both weighted and transition methods.

In addition to the peak around 1, a peak ranging from 2-3 was observed in the coefficient distribution in the Mycobacteriales dataset (Fig. 4A). This peak mainly comprised gene pairs of flagellar COGs belonging to categories N and/or U (Supplementary Fig. S6). We noticed that the differences in the number of presence/absence state changes across the phylogenetic tree (i.e., t_1_ and t_2_ in SEV) were greater for flagellar COG pairs detected only by the transition methods than the other pairs, except for two that were only detected by ASA (Fig. 4B and Supplementary Figs. S7-S9). The asymmetry in t_1_ and t_2_ led to an uneven distribution of the fourteen COGs found only in the pairs detected by the transition methods and their co-evolved partners (Supplementary Fig S10 and Fig. 4C). Therefore, it is difficult to identify them using weighted methods. The reason why the fourteen genes were independently gained or lost from the other flagellar proteins is unclear, but they can have another function or work with other flagellar proteins that have not yet been defined in the COG. It is also possible that some of the homologous genes of the fourteen COGs have evolved to dissimilar amino acid sequences because of different evolutionary pressures, preventing them from being classified into the same COGs.

We also examined the effect of varying TPR. At the highest TPR (0.9), the peaks of the COG category J in transition methods were not observed in all datasets (Supplementary Figure S11). Alternatively, there is a peak of the COG category N in the Pseudomonadales datasets at TPR=0.9. Unlike the peak comprising COG category N/U in the Mycobacteriales at lower TPRs, the differences of t1 and t2 of the pairs consisting of this peak were similar between the pair only detected by transition methods and the others (Supplementary Figure S12). Moreover, these pairs were found by weighted methods at lower TPRs, resulting in the disappearance of this peak. On the contrary, decreasing the TPR threshold from 0.7 does not substantially affect the coefficient distribution across the COG category in transition methods (Supplementary Figures S13 and S14). Consequently, the detection of highly conserved pairs and unevenly distributed pairs is a characteristic of transition methods.

### 3.3 Weighted methods can identify the gain/loss order of the gene pairs

Although the transition methods found more positive pairs than any other weighted methods, there are pairs detected only by weighted methods and not by transition methods (Supplementary Tables S5-S7). Considering that transition methods require simultaneous gains and losses of OGs, they can miss pairs evolving through the intermediate state, where only one OG is present. In contrast, weighted methods do not matter if the OGs have evolved simultaneously. To investigate this, we examined the changes that led to the current states of the pair profiles. Specifically, we divided the phylogenetic tree into subtrees based on the reconstructed ancestral state of the pair profiles, as in the first step of ASA, and then checked the state change between the root of each subtree and its ancestral node. As expected, the transitions from/to the intermediate state were more frequently observed at the root of the subtrees in pairs detected only by weighted pairs than in those detected only by transition methods at all evaluated TPRs (Supplementary Figure S15).

We further investigated which intermediate state is preferred in each pair. If one of the intermediate states appears frequently across the phylogenetic tree and the other does not, there should be a strong evolutionary constraint on the order of OG gain/loss. Intriguingly, most of state changes in the pair detected only by the weighted method in the Pseudomonadales dataset were biased to one direction (Fig. 5). The same trends were observed in the other two datasets and at all examined TPRs, but they are not as clear as in the Pseudomonadales dataset (Fig. 5 and Supplementary Figure S16), partially because a smaller number of pairs were only detected by the weighted methods.

**Figure 5.**
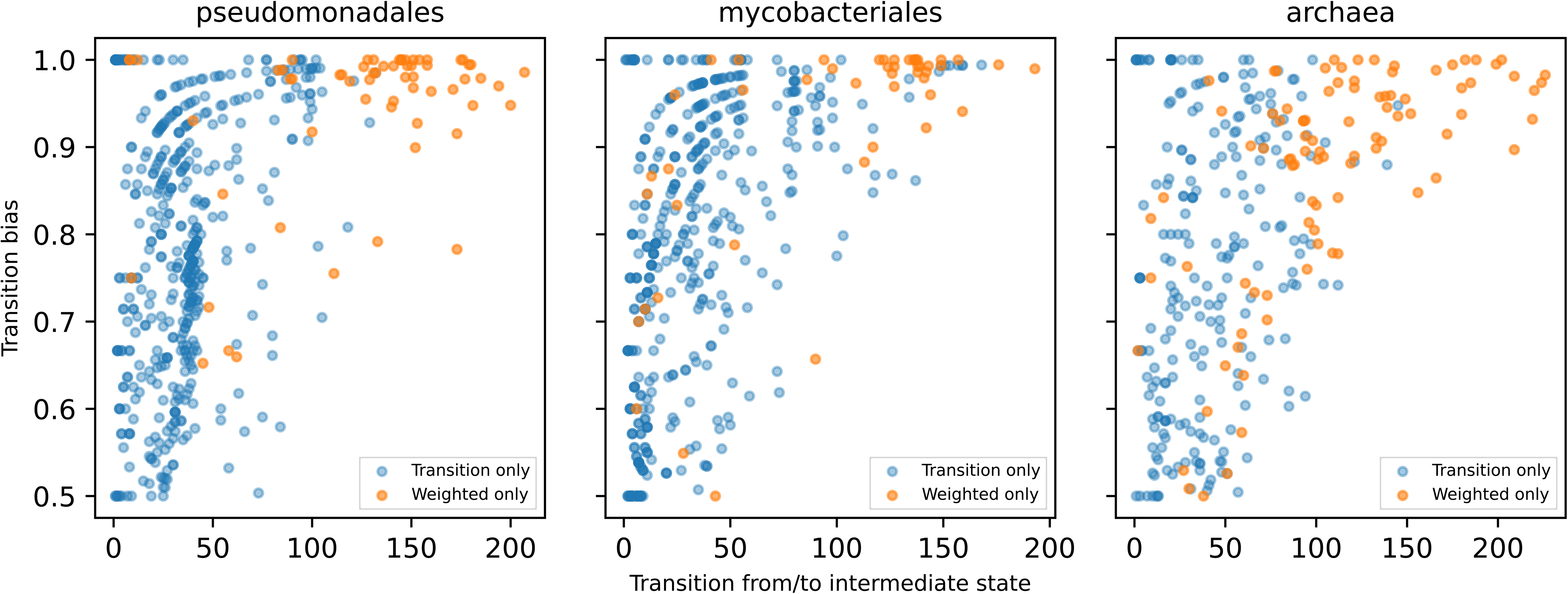
Characteristic of the weighted methods. Comparison of the numbers of changes from/to the intermediate state of the co-evolved gene pairs only detected by the transition method (blue) and weighted methods (orange) at a True Positive Rate (TPR) =0.7. The number of changes was counted from the ancestral state reconstruction result of the co-evolved gene pairs. The numbers of the changes from/to the intermediate state are indicated on the x-axis, and the biases to the changes to one of two intermediate states are indicated on the y-axis. The transition bias was calculated by dividing the larger number of changes from/to the intermediate state of pair genes by their sum. The plots at the other TPRs are shown in Supplementary Figure S16.

Among the three datasets, the COGs for Type I restriction-modification (RM) systems were commonly found in the lists of co-evolved pairs with fixed-order evolution at various TPRs (see Supplementary Tables S8-S10 available at Zenodo). The genes for methyltransferase (MTase; COG0286) and specificity (S) subunit (COG0732) were obtained before the acquisition of restriction endonucleases (REases; COG4096), or alternatively, REases may have been lost earlier than MTase and the S subunit genes. In prokaryotes, Type I restriction-modification (RM) system works as an immunity against foreign DNA, such as phages and plasmids (41). REase digests foreign, non-methylated DNA, but does not digest host DNA methylated by MTase. The S subunit is a characteristic of Type I RM system and determines the specificity of REase and MTase. As REase without MTase and the S subunit may have caused host DNA digestion, it is reasonable to avoid the state with REase but without MTase and the S subunit during evolution.

Similarly, a state with antitoxin alone is permitted for the toxin-antitoxin (TA) system in Pseudomonadales and Mycobacteriales. The toxin is neutralized by forming a complex with the antitoxin when a bacterial cell is not infected by the phage. Once a bacterial cell is infected, the toxin is activated, leading to cell death to prevent phage production (42). Therefore, the state with a toxin alone is harmful to the cell, and it is reasonable that the TA system has evolved through the other state.

The examples mentioned above are only part of the co-evolved pairs with a fixed intermediate state, and their biological interpretation remains to be resolved in future studies.

### 3.4 Runtime performance in simulated datasets

The runtimes for each method were evaluated using two groups of simulated datasets. One group contained 1, 000 genomes with varying numbers of genes ranging from 2, 000 to 64, 000. As CWA and ASA have to parse all nodes in the tree for all combinations of gene pairs, they require a longer time than the other methods, and their runtime increases with the square of the number of OGs (Fig. 6A). The RLE runtime also increased in the same order, but the time required for the one pair comparison was much shorter (Fig. 6A). Compared to these three methods, naive, cotransitions, and SEV performed very quickly, as they can leverage matrix multiplication for all-vs-all gene comparisons (Fig. 6A). Calculation with GPU further accelerates these methods; a dataset with 64, 000 genes (∼2 billion pair comparisons) was calculated within 20 min.

**Figure 6.**
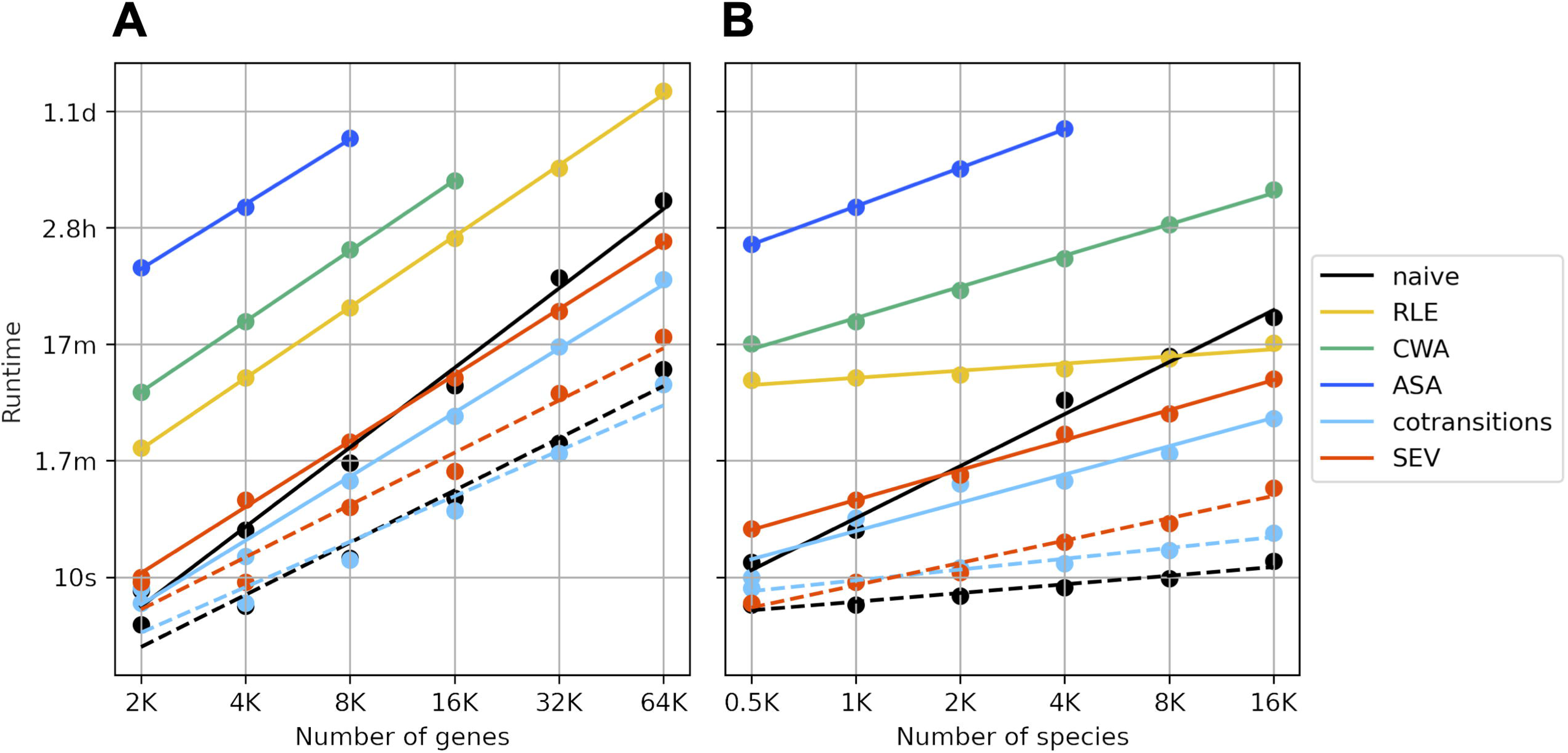
Runtimes comparison among phylogenetic profiling methods with (A) 1, 000 genomes and a variety number of genes and (B) 4, 000 genes and a variety number of genomes. The runtimes of naive, RLE, CWA, ASA, cotransition, and SEV are indicated by the black, yellow, green, blue, sky blue, and red circles, respectively, and the lines indicate the regression curve of the runtimes. The runtimes with the GPU are connected by the dotted lines.

The other groups of simulated data included varying numbers of genomes, from 500 to 16, 000 and 4, 000 genes. Again, ASA was the most time-consuming method, indicating that it is difficult to apply to a large-scale dataset (Fig. 6B). The runtime increase was slower than linear with the number of genomes, except for ASA and naive without GPU.

As the runtime of each method consistently increased with the number of genomes and genes, we inferred the runtime with multiple regression to predict the runtime of arbitrary numbers of genomes and genes. The regressions fitted the data well, with R^2^ > 0.95 (Supplementary Table S11). Transition methods with GPU were predicted to conduct comparisons of all gene pairs in the datasets as large as the latest OrthoDB (v12) and GTDB (Release 220) within an hour (Supplementary Table S11), suggesting their high scalability.

### 3.5 Performance comparison among ASA, SEV and EvoWeaver

At the final stage of manuscript preparation, a new tool named EvoWeaver has been published (32). EvoWeaver implements twelve methods to detect coevolution signals in OG pairs and integrates the results by a machine learning approach. Four of twelve methods belong to phylogenetic profiling that capture different evolutionary perspectives from the distribution of presence/absence (P/A) or gain/loss (G/L) of paired OGs; “P/A Jaccard” and “G/L MI” are similar approaches to CWA and SEV, respectively. “P/A Overlap” uses branch lengths to detect coevolution signals, rather than the number of genomes. Finally, “G/L Distance” attaches importance to the shortness of the distance (branch length) between the gain/loss events of paired OG. Statistical significance in these methods is evaluated by combining the respective measures of each method and the p-value derived from the permutation test.

To compare our methods with phylogenetic profiling methods in EvoWeaver, we analyzed three prokaryotic datasets as in the previous section. As a result, G/L Distance showed the best or second-best PRAUC in all datasets (Fig. 7A and Supplementary Figure S17). On the other hand, the other three methods in EvoWeaver showed modest performance, and we therefore highlight G/L Distance hereafter. Intuitively, G/L Distance can detect both OG pairs where two genes are simultaneously gained/lost (i.e., distance is 0) and those where the gain/loss of two genes occurs at different timings. Thus, G/L Distance seems to have both the strengths of weighted and transition methods. In fact, G/L Distance detected most or the second most positive pairs at TPR=0.7-0.9, except in the Mycobacteriales datasets at TPR=0.9 (Fig. 7B and Supplementary Figure S18). However, the number of positive pairs detected by G/L Distance were outperformed by SEV at the lower TPRs (=0.5 and 0.6) and the set of positive pairs detected by SEV and G/L Distance is almost overlapped (Supplementary Figure S19), indicating that the pairs detected only by G/L Distance at higher TPRs are not only gained/lost at slightly different timings but also simultaneously in the evolution.

**Figure 7.**
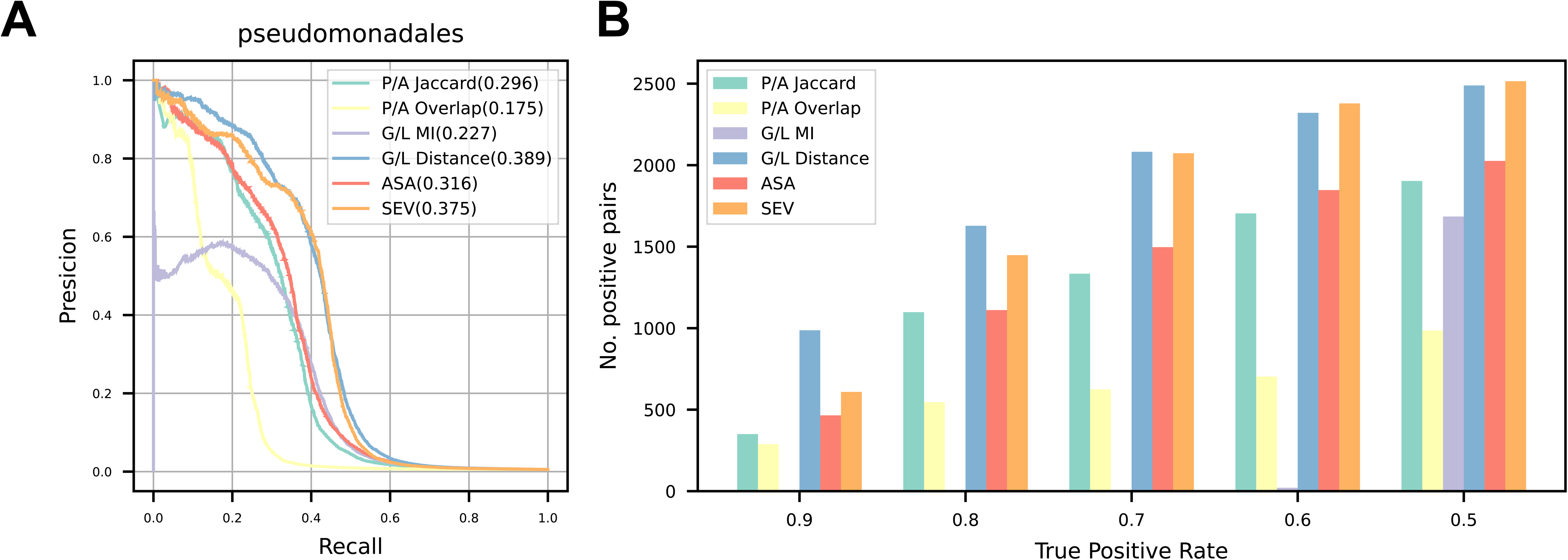
Performance comparison of phylogenetic profiling methods in this study and EvoWeaver using the Pseudomonadales datasets. (A) Precision-Recall (PR) curves of each method. Areas under the PR curve (PRAUCs) are shown in parentheses. (B) The number of detected positive pairs of each method at various true positive rate (TPR) thresholds. The comparison using the other datasets are shown in Supplementary Figures S17 and S18.

One of the drawbacks of G/L Distance is its runtime performance. Although the most time-consuming parts of G/L Distance are implemented in the C language, the permutation test is computationally burdensome. Therefore, it is difficult for G/L Distance to analyze all OG pairs in a large dataset containing thousands or tens of thousands of genomes and OGs (Supplementary Table S11). It is also worth noting that the runtimes for datasets with more than 32, 000 OGs could not be measured due to memory limitations. This indicates that G/L Distance shows limited memory efficiency during parallel execution, which constrains its applicability to very large datasets.

In summary, G/L Distance is better than or competitive to ASA and SEV in terms of precision, but SEV is much more scalable than G/L Distance. Combining CORGIAS and EvoWeaver could improve the detection of functionally related gene sets in the future.

## 4. Discussion

In this study, we developed two novel methods for phylogenetic profiling: ASA and SEV. Considering the ancestral state and gene gain/loss events were handled more properly than in the previous method, their results are robust to branch rotation. In addition, ASA showed better recall than similar methods in the validation datasets. SEV also usually detected more co-evolved pairs than its predecessor, cotransitions, although the difference was small. Furthermore, SEV and cotransitions implemented with matrix manipulation showed high scalability, which enables us to investigate the latest large-scale databases. ASA, SEV, and the other phylogenetic profiling methods compared in this study are freely available in a Python package CORGIAS (CORelated Genes Identifier by Considering Ancestral State; https://github.com/ynishimuraLv/corgias.git).

Although the weighted methods, including ASA, were outperformed by the transition methods in the number of detected positive pairs, they are able to capture the co-evolved pairs that evolved through the intermediate states. Unexpectedly, in the Pseudomonadales dataset, most of the pairs detected only by the weighted methods have evolved through only one of the two possible intermediate states. The reason this trend is remarkable in the Pseudomonadales dataset alone is unclear, but the largest number of genomes among the three datasets can affect it because the trend gets weaker as the number of genomes decreases. The accuracy and granularity of ortholog clustering (e.g., COG classification is regarded as coarser than the KEGG ortholog) and other possible factors should be investigated in future studies. To further investigate the characteristics of OG pairs detected only by weighted methods, ASA must be accelerated, as it is the slowest method in this study. As EvoWeaver, implementing ASA with faster programming languages is one of the possible solutions but improving the algorithm is also possible. From the perspective of evolutionary biology, the interpretation of the order of gene gain/loss is another interesting topic (43).

Phylogenetic profiling depends on the species tree, intrinsically including uncertainty. Although the phylogenetic trees used in this study were not prepared by the best effort, ASA and SEV showed better or comparable performance to previous and recently published methods, suggesting their robustness and potential advantage in tree uncertainty. In addition to tree uncertainty, ASA and SEV depend on how the ancestral states of the profile are reconstructed. The ancestral state reconstruction methods that we adopted were quick but simple and ignored the intrinsic complexity of evolution. For example, the gain/loss rate heterogeneity among genes and lineages, (44–46), recombination, and horizontal gene transfers (47) can mislead inferences under the simple evolutionary model. Although ancestral state reconstruction methods considering complex evolutionary backgrounds and phylogenetic reconciliation that can capture horizontal transfer are computationally expensive and contain many uncertainties (48), it is worth investigating whether they, together with the more accurate species tree, can improve ASA and SEV. On the other hand, the recent study indicates ancestral state reconstruction based on a simple model can detect more HGTs than phylogenetic reconciliation (49). Actually, the correlated pairs detected by ASA and SEV include those for anti-phage systems such as the RM system, the TA system, and the CRISPR-Cas system and those encoded in prophage regions that frequently transfer among distantly related prokaryotes (50). The influence of HGTs on ASA and SEV (and other methods utilizing ancestral state) should be evaluated in future studies.

The phylogenetic profiling methods developed in this study can be applied to any binary dataset and are not limited to gene absence/presence data. They can be useful in finding genotype-phenotype associations when gene profiles and binary trait data are used, as in a trait-based approach (34, 51). Improvements to allow for the handling of categorical and quantitative data, including copy number or OGs, are a possible direction for future studies. As the methods in this study can detect only paired relationships, detecting the relationship among more than three genes (24) and/or considering the global metric (16, 52) is another direction. Machine learning approach may be a promising approach in these directions (43, 53).

In conclusion, CORGIAS is an accurate and scalable phylogenetic profiling method. Our method is powerful for narrowing down OGs with a target function from a number of unannotated OGs encoded in microbial dark matter. Applying the candidate OGs identified by CORGIAS to other bioinformatics methods, such as those using synteny (54, 55), gene tree (11), co-expression(56, 57), protein structure (58, 59), and a combination of them (32, 38), further provides clues for functional prediction.

## Data availability

The implementation of CORGIAS is freely available at (https://github.com/ynishimuraLv/corgias(doi: 10.5281/zenodo.15583905). The genome data used in this study can be found in GTDB release 214. The scores of the COG pair can be downloaded from STRING database Version 12.0. The codes and intermediate files to reproduce all the results in this paper are deposited at https://github.com/ynishimuraLv/corgias_data and Zenodo (doi: 10.5281/zenodo.15496141).

## Supplementary Data

Supplementary data are provided as separate files.

## Author Contribution

Yuki Nishimura: Conceptualization, Data curation, Formal analysis, Investigation, Methodology, Software, Validation, Visualization, Writing – original draft, Writing – review & editing. Kimiho Omae: Writing – review & editing. Kento Tominaga: Writing – review & editing. Wataru Iwasaki, Conceptualization, Funding acquisition, Writing – review & editing.

## Acknowledgment

The authors thank Dr. Wataru Iwasaki’s lab members for the fruitful discussion.

## References

1. Brown, C.T., Hug, L.A., Thomas, B.C., Sharon, I., Castelle, C.J., Singh, A., Wilkins, M.J., Wrighton, K.C., Williams, K.H. and Banfield, J.F. (2015) Unusual biology across a group comprising more than 15% of domain Bacteria. Nature, 523, 208–211.

2. Hug, L.A., Baker, B.J., Anantharaman, K., Brown, C.T., Probst, A.J., Castelle, C.J., Butterfield, C.N., Hernsdorf, A.W., Amano, Y., Ise, K., et al. (2016) A new view of the tree of life. Nat Microbiol, 1, 16048.

3. Nayfach, S., Roux, S., Seshadri, R., Udwary, D., Varghese, N., Schulz, F., Wu, D., Paez-Espino, D., Chen, I.-M., Huntemann, M., et al. (2021) A genomic catalog of Earth’s microbiomes. Nat. Biotechnol., 39, 499–509.

4. Almeida, A., Nayfach, S., Boland, M., Strozzi, F., Beracochea, M., Shi, Z.J., Pollard, K.S., Sakharova, E., Parks, D.H., Hugenholtz, P., et al. (2021) A unified catalog of 204, 938 reference genomes from the human gut microbiome. Nat. Biotechnol., 39, 105–114.

5. Baltoumas, F.A., Karatzas, E., Liu, S., Ovchinnikov, S., Sofianatos, Y., Chen, I.-M., Kyrpides, N.C. and Pavlopoulos, G.A. (2023) NMPFamsDB: a database of novel protein families from microbial metagenomes and metatranscriptomes. Nucleic Acids Res., 52, D502–D512.

6. Río, Á.R. del, Giner-Lamia, J., Cantalapiedra, C.P., Botas, J., Deng, Z., Hernández-Plaza, A., Munar-Palmer, M., Santamaría-Hernando, S., Rodríguez-Herva, J.J., Ruscheweyh, H.-J., et al. (2024) Functional and evolutionary significance of unknown genes from uncultivated taxa. Nature, 626, 377–384.

7. Castelle, C.J., Brown, C.T., Anantharaman, K., Probst, A.J., Huang, R.H. and Banfield, J.F. (2018) Biosynthetic capacity, metabolic variety and unusual biology in the CPR and DPANN radiations. Nat Rev Microbiol, 16, 629–645.

8. Pellegrini, M., Marcotte, E.M., Thompson, M.J., Eisenberg, D. and Yeates, T.O. (1999) Assigning protein functions by comparative genome analysis: Protein phylogenetic profiles. Proc. Natl. Acad. Sci., 96, 4285–4288.

9. Tabach, Y., Billi, A.C., Hayes, G.D., Newman, M.A., Zuk, O., Gabel, H., Kamath, R., Yacoby, K., Chapman, B., Garcia, S.M., et al. (2013) Identification of small RNA pathway genes using patterns of phylogenetic conservation and divergence. Nature, 493, 694–698.

10. Dey, G., Jaimovich, A., Collins, S.R., Seki, A. and Meyer, T. (2015) Systematic Discovery of Human Gene Function and Principles of Modular Organization through Phylogenetic Profiling. Cell Rep., 10, 993–1006.

11. Moi, D., Kilchoer, L., Aguilar, P.S. and Dessimoz, C. (2020) Scalable phylogenetic profiling using MinHash uncovers likely eukaryotic sexual reproduction genes. PLoS Comput. Biol., 16, e1007553.

12. Dembech, E., Malatesta, M., Rito, C.D., Mori, G., Cavazzini, D., Secchi, A., Morandin, F. and Percudani, R. (2023) Identification of hidden associations among eukaryotic genes through statistical analysis of coevolutionary transitions. Proc. Natl. Acad. Sci., 120, e2218329120.

13. Li, Y., Ma, B., Hua, K., Gong, H., He, R., Luo, R., Bi, D., Zhou, R., Langford, P.R. and Jin, H. (2023) PPNet: Identifying Functional Association Networks by Phylogenetic Profiling of Prokaryotic Genomes. Microbiol Spectr, 11, e03871–22.

14. Fukunaga, T., Ogawa, T., Iwasaki, W. and Sonoike, K. (2024) Phylogenetic Profiling Analysis of the Phycobilisome Revealed a Novel State-Transition Regulator Gene in Synechocystis sp. PCC 6803. Plant Cell Physiol., 65, 1450–1460.

15. Whelan, F.J., Rusilowicz, M. and McInerney, J.O. (2020) Coinfinder: detecting significant associations and dissociations in pangenomes. Microb. Genom., 6, e000338.

16. Fukunaga, T. and Iwasaki, W. (2022) Inverse Potts model improves accuracy of phylogenetic profiling. Bioinformatics, 38, 1794–1800.

17. Morett, E., Korbel, J.O., Rajan, E., Saab-Rincon, G., Olvera, L., Olvera, M., Schmidt, S., Snel, B. and Bork, P. (2003) Systematic discovery of analogous enzymes in thiamin biosynthesis. Nat. Biotechnol., 21, 790–795.

18. Kumagai, Y., Yoshizawa, S., Nakajima, Y., Watanabe, M., Fukunaga, T., Ogura, Y., Hayashi, T., Oshima, K., Hattori, M., Ikeuchi, M., et al. (2018) Solar-panel and parasol strategies shape the proteorhodopsin distribution pattern in marine Flavobacteriia. Isme J, 12, 1329–1343.

19. Gavriilidou, A., Paulitz, E., Resl, C., Ziemert, N., Kupczok, A. and Baumdicker, F. (2024) Goldfinder: Unraveling Networks of Gene Co-occurrence and Avoidance in Bacterial Pangenomes. bioRxiv, 10.1101/2024.04.29.591652.

20. Tominaga, K., Ozaki, S., Sato, S., Katayama, T., Nishimura, Y., Omae, K. and Iwasaki, W. (2024) Frequent nonhomologous replacement of replicative helicase loaders by viruses in Vibrionaceae. Proc. Natl. Acad. Sci. United States Am., 121, e2317954121.

21. Pagel, M. (1994) Detecting correlated evolution on phylogenies: a general method for the comparative analysis of discrete characters. Proc. R. Soc. Lond. Ser. B: Biol. Sci., 255, 37–45.

22. Cohen, O., Ashkenazy, H., Karin, E.L., Burstein, D. and Pupko, T. (2013) CoPAP: Coevolution of Presence–Absence Patterns. Nucleic Acids Res., 41, W232–W237.

23. Li, Y., Calvo, S.E., Gutman, R., Liu, J.S. and Mootha, V.K. (2014) Expansion of Biological Pathways Based on Evolutionary Inference. Cell, 158, 213–225.

24. Liu, C., Kenney, T., Beiko, R.G. and Gu, H. (2022) The Community Coevolution Model with Application to the Study of Evolutionary Relationships between Genes Based on Phylogenetic Profiles. Systematic Biol, 10.1093/sysbio/syac052.

25. Parks, D.H., Chuvochina, M., Waite, D.W., Rinke, C., Skarshewski, A., Chaumeil, P.-A. and Hugenholtz, P. (2018) A standardized bacterial taxonomy based on genome phylogeny substantially revises the tree of life. Nat Biotechnol, 36, 996–1004.

26. Rinke, C., Chuvochina, M., Mussig, A.J., Chaumeil, P.-A., Davín, A.A., Waite, D.W., Whitman, W.B., Parks, D.H. and Hugenholtz, P. (2021) A standardized archaeal taxonomy for the Genome Taxonomy Database. Nat Microbiol, 6, 946–959.

27. Mendler, K., Chen, H., Parks, D.H., Lobb, B., Hug, L.A. and Doxey, A.C. (2019) AnnoTree: visualization and exploration of a functionally annotated microbial tree of life. Nucleic Acids Res., 47, 4442–4448.

28. Cokus, S., Mizutani, S. and Pellegrini, M. (2007) An improved method for identifying functionally linked proteins using phylogenetic profiles. Bmc Bioinformatics, 8, S7–S7.

29. Tremblay, B.J.-M., Lobb, B. and Doxey, A.C. (2021) PhyloCorrelate: inferring bacterial gene-gene functional associations through large-scale phylogenetic profiling. Bioinform Oxf Engl, 37, 17–22.

30. Mering, C. von, Huynen, M., Jaeggi, D., Schmidt, S., Bork, P. and Snel, B. (2003) STRING: a database of predicted functional associations between proteins. Nucleic Acids Res., 31, 258–261.

31. Ruano-Rubio, V., Poch, O. and Thompson, J.D. (2009) Comparison of eukaryotic phylogenetic profiling approaches using species tree aware methods. BMC Bioinform., 10, 383–383.

32. Lakshman, A.H. and Wright, E.S. (2025) EvoWeaver: large-scale prediction of gene functional associations from coevolutionary signals. Nat. Commun., 16, 3878.

33. Ishikawa, S.A., Zhukova, A., Iwasaki, W. and Gascuel, O. (2019) A Fast Likelihood Method to Reconstruct and Visualize Ancestral Scenarios. Mol. Biol. Evol., 36, 2069–2085.

34. Collins, C. and Didelot, X. (2018) A phylogenetic method to perform genome-wide association studies in microbes that accounts for population structure and recombination. PLoS Comput. Biol., 14, e1005958.

35. Parks, D.H., Chuvochina, M., Rinke, C., Mussig, A.J., Chaumeil, P.-A. and Hugenholtz, P. (2021) GTDB: an ongoing census of bacterial and archaeal diversity through a phylogenetically consistent, rank normalized and complete genome-based taxonomy. Nucleic Acids Res, 50, gkab776-.

36. Chklovski, A., Parks, D.H., Woodcroft, B.J. and Tyson, G.W. (2023) CheckM2: a rapid, scalable and accurate tool for assessing microbial genome quality using machine learning. Nat. Methods, 20, 1203–1212.

37. Galperin, M.Y., Wolf, Y.I., Makarova, K.S., Alvarez, R.V., Landsman, D. and Koonin, E.V. (2020) COG database update: focus on microbial diversity, model organisms, and widespread pathogens. Nucleic Acids Res., 49, D274–D281.

38. Szklarczyk, D., Kirsch, R., Koutrouli, M., Nastou, K., Mehryary, F., Hachilif, R., Gable, A.L., Fang, T., Doncheva, N.T., Pyysalo, S., et al. (2022) The STRING database in 2023: protein–protein association networks and functional enrichment analyses for any sequenced genome of interest. Nucleic Acids Res., 51, D638–D646.

39. Revell, L.J. (2012) phytools: an R package for phylogenetic comparative biology (and other things). Methods Ecol. Evol., 3, 217–223.

40. Ramulu, H.G., Groussin, M., Talla, E., Planel, R., Daubin, V. and Brochier-Armanet, C. (2014) Ribosomal proteins: Toward a next generation standard for prokaryotic systematics? Mol. Phylogenetics Evol., 75, 103–117.

41. Loenen, W.A.M., Dryden, D.T.F., Raleigh, E.A. and Wilson, G.G. (2014) Type I restriction enzymes and their relatives. Nucleic Acids Res., 42, 20–44.

42. Sberro, H., Leavitt, A., Kiro, R., Koh, E., Peleg, Y., Qimron, U. and Sorek, R. (2013) Discovery of Functional Toxin/Antitoxin Systems in Bacteria by Shotgun Cloning. Mol Cell, 50, 136–148.

43. Konno, N. and Iwasaki, W. (2023) Machine learning enables prediction of metabolic system evolution in bacteria. Sci. Adv., 9, eadc9130.

44. Fukunaga, T. and Iwasaki, W. (2021) Mirage: estimation of ancestral gene-copy numbers by considering different evolutionary patterns among gene families. Bioinform. Adv., 1, vbab014.

45. Fukunaga, T. and Iwasaki, W. (2022) Mirage 2.0: fast and memory-efficient reconstruction of gene-content evolution considering heterogeneous evolutionary patterns among gene families. Bioinformatics, 38, 4039–4041.

46. Yamanouchi, S., Fukunaga, T. and Iwasaki, W. (2024) CoLaML: Inferring latent evolutionary modes from heterogeneous gene content. bioRxiv, 10.1101/2024.12.02.626417.

47. Dagan, T. and Martin, W. (2007) Ancestral genome sizes specify the minimum rate of lateral gene transfer during prokaryote evolution. Proc. Natl. Acad. Sci., 104, 870–875.

48. Williams, T.A., Davin, A.A., Szánthó, L.L., Stamatakis, A., Wahl, N.A., Woodcroft, B.J., Soo, R.M., Eme, L., Sheridan, P.O., Gubry-Rangin, C., et al. (2024) Phylogenetic reconciliation: making the most of genomes to understand microbial ecology and evolution. ISME J., 18, wrae129.

49. Mishra, S. and Lercher, M.J. (2025) Horizontal Gene Transfer Inference: Gene Presence–Absence Outperforms Gene Trees. Mol. Biol. Evol., 42, msaf166.

50. Koonin, E.V., Makarova, K.S. and Wolf, Y.I. (2016) Evolutionary Genomics of Defense Systems in Archaea and Bacteria. Annu Rev Microbiol, 71, 1–29.

51. Yang, Y. and Jiang, X. (2023) Evolink: a phylogenetic approach for rapid identification of genotype–phenotype associations in large-scale microbial multispecies data. Bioinformatics, 39, btad215.

52. Kim, P.-J. and Price, N.D. (2011) Genetic Co-Occurrence Network across Sequenced Microbes. PLoS Comput. Biol., 7, e1002340.

53. Beavan, A.J.S., Domingo-Sananes, M.R. and McInerney, J.O. (2024) Contingency, repeatability, and predictability in the evolution of a prokaryotic pangenome. Proc. Natl. Acad. Sci., 121, e2304934120.

54. Shmakov, S.A., Faure, G., Makarova, K.S., Wolf, Y.I., Severinov, K.V. and Koonin, E.V. (2019) Systematic prediction of functionally linked genes in bacterial and archaeal genomes. Nat. Protoc., 14, 3013–3031.

55. Botas, J., Río, Á.R. del, Giner-Lamia, J. and Huerta-Cepas, J. (2022) GeCoViz: genomic context visualisation of prokaryotic genes from a functional and evolutionary perspective. Nucleic Acids Res, 50, W352–W357.

56. Dam, S. van, Võsa, U., Graaf, A. van der, Franke, L. and Magalhães, J.P. de (2017) Gene co-expression analysis for functional classification and gene–disease predictions. Brief. Bioinform., 19, 575–592.

57. Chang, L.-Y., Lee, M.-Z., Wu, Y., Lee, W.-K., Ma, C.-L., Chang, J.-M., Chen, C.-W., Huang, T.-C., Lee, C.-H., Lee, J.-C., et al. (2023) Gene set correlation enrichment analysis for interpreting and annotating gene expression profiles. Nucleic Acids Res., 52, e17–e17.

58. Jumper, J., Evans, R., Pritzel, A., Green, T., Figurnov, M., Ronneberger, O., Tunyasuvunakool, K., Bates, R., Žídek, A., Potapenko, A., et al. (2021) Highly accurate protein structure prediction with AlphaFold. Nature, 596, 583–589.

59. Abramson, J., Adler, J., Dunger, J., Evans, R., Green, T., Pritzel, A., Ronneberger, O., Willmore, L., Ballard, A.J., Bambrick, J., et al. (2024) Accurate structure prediction of biomolecular interactions with AlphaFold 3. Nature, 630, 493–500.

