## Supplementary Figures S1-19 for "CORGIAS: identifying correlated gene pairs by considering evolutionary history in a large-scale prokaryotic genome dataset"

**Supplementary Figure S1.** STRING score distribution. Distributions of COG pairs's scores in the three prokaryotic datasets. Left, distributions of all scored pairs. Right, score distributions of top-ranking pairs before the True Positive rates fell under 50%. The top-ranking pairs were accumulated across the six phylogenetic profiling methods (naive, RLE, CWA, ASA, cotransitions, and SEV) analyzed in this study. Red dot lines indicate a score of 900, the threshold of functional correlation in this study.

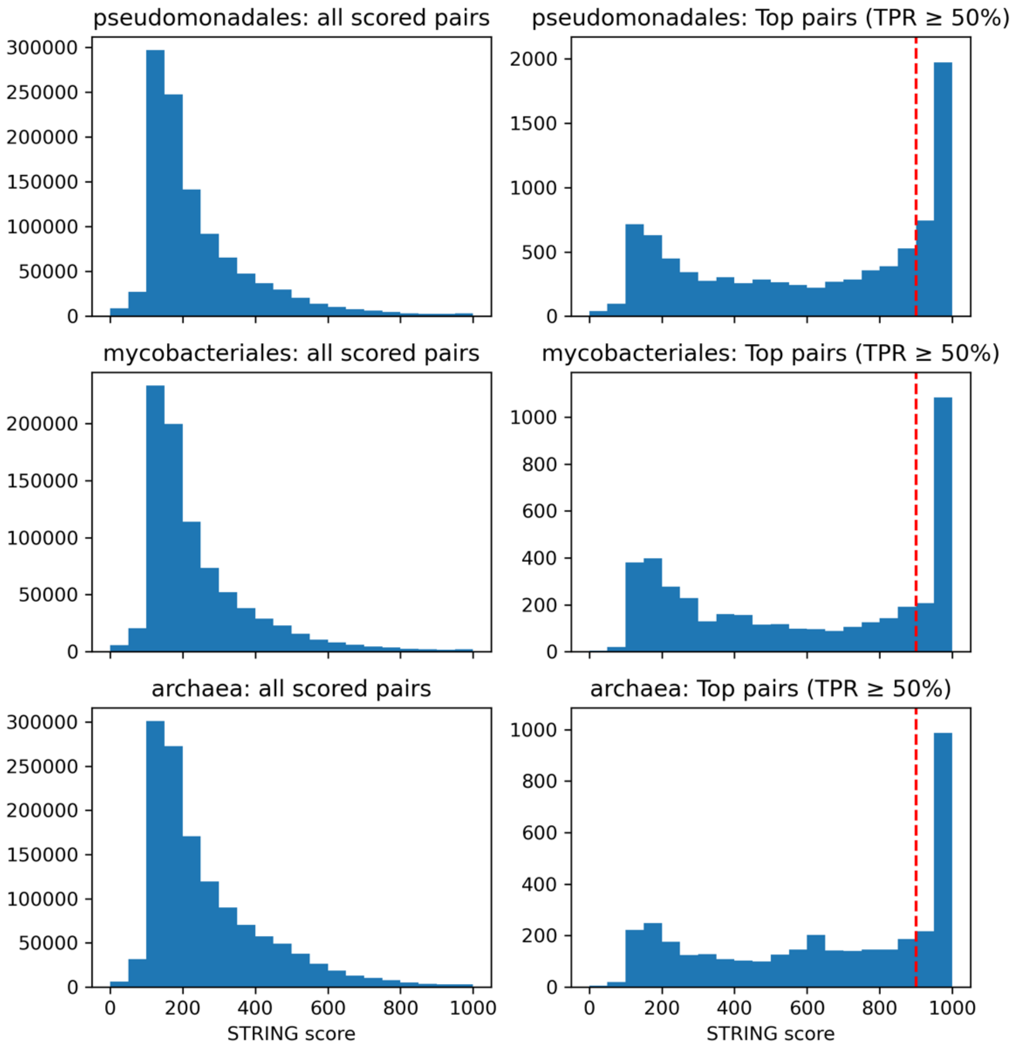

**Supplementary Figure S2.** The number of unique pairs detected by each phylogenetic profiling method. The number of positive pairs only detected by the tagged methods but not by the other methods is shown at various True Positive rates.

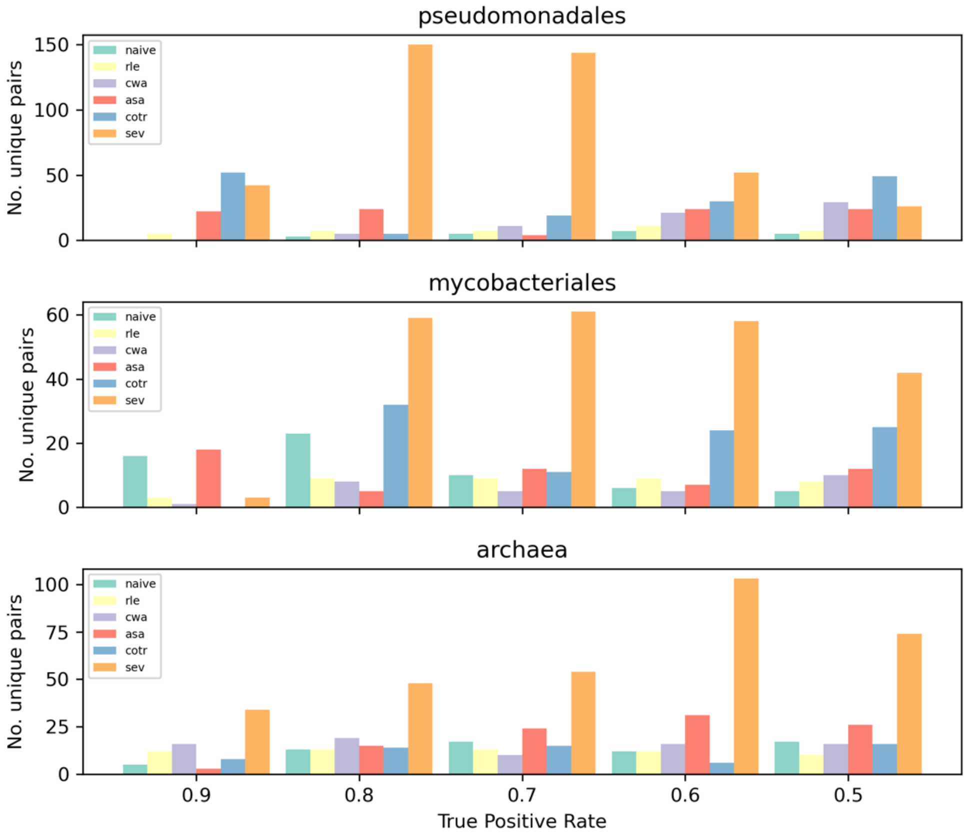

**Supplementary Figure S3.** Comparison of cotransitions and SEV. The evolCCM coefficient distribution of pairs detected by cotransitions (cotr, blue) but not SEV (orange), and vice versa, at various true positive rates.

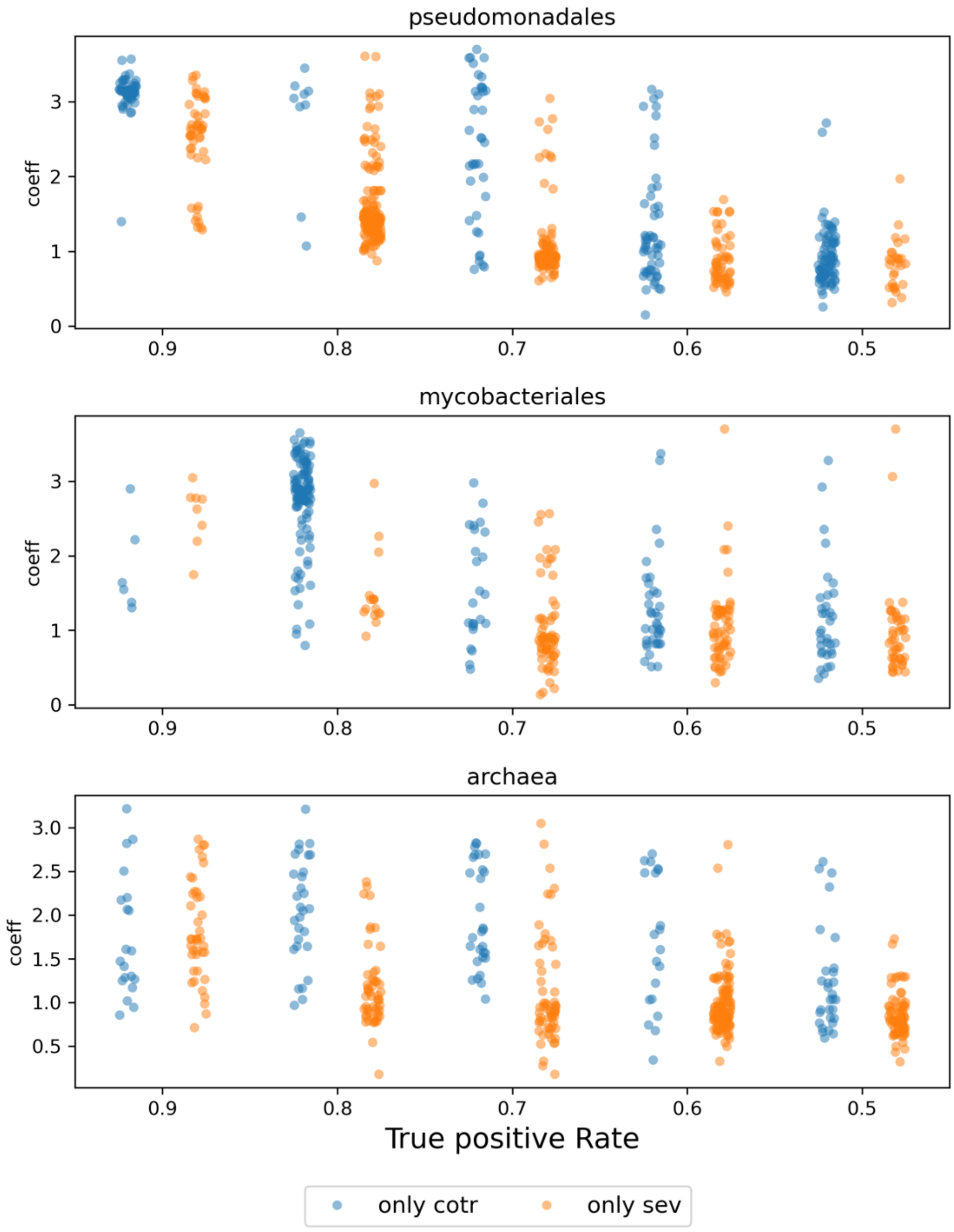

**Supplementary Figure S4.** Comparison of CWA and ASA. The evolCCM coefficient distribution of pairs detected by CWA (blue) but not ASA (orange), and vice versa, at various true positive rates.

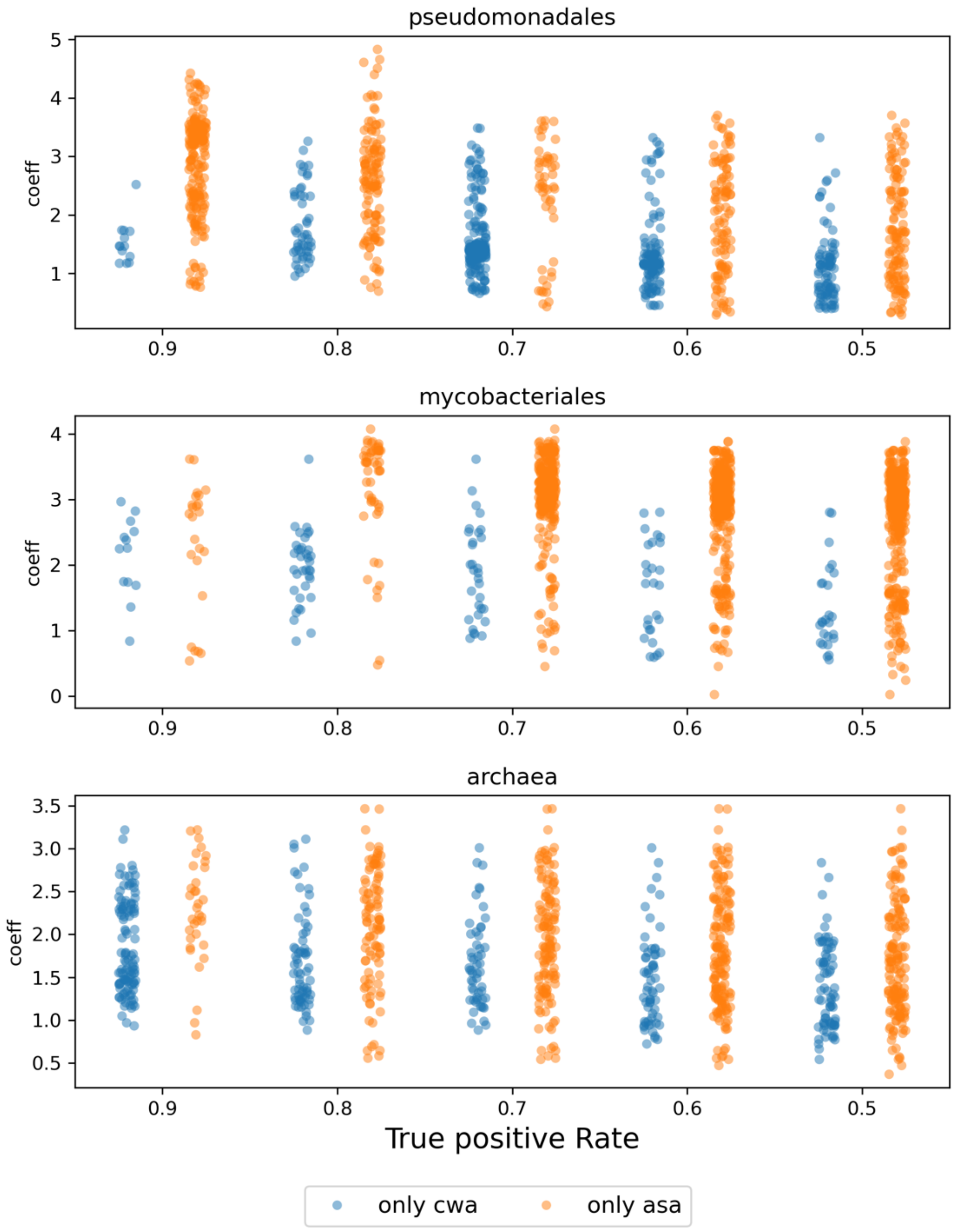

**Supplementary Figure S5.** Comparison of RLE and ASA. The evolCCM coefficient distribution of pairs detected by RLE (blue) but not ASA (orange), and vice versa, at various true positive rates.

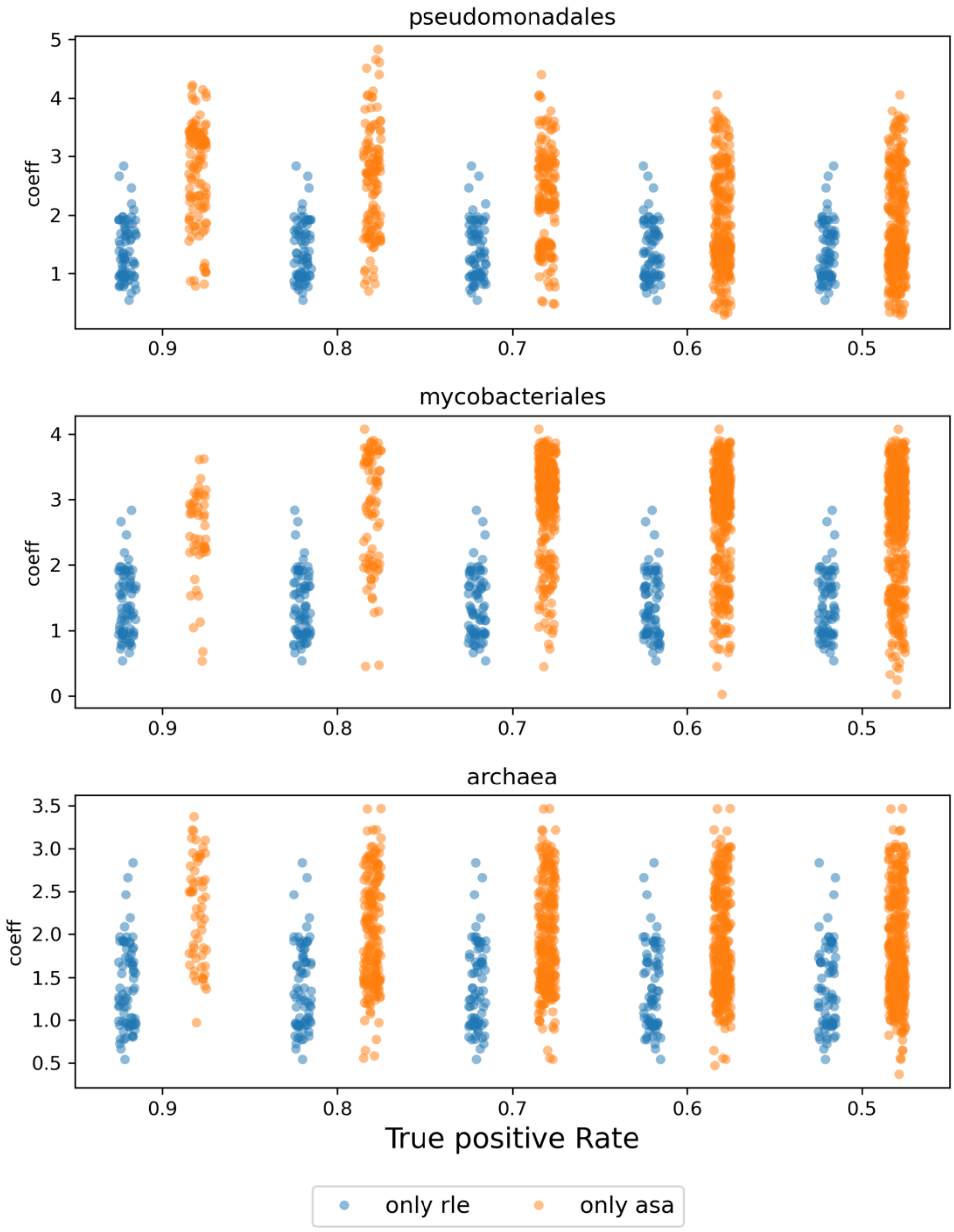

**Supplementary Figure S6.** The distribution of coefficients of the detected positive pairs only detected by transition methods (green), weighted methods (orange), and both (blue) at a True Positive Rate (TPR) = 0.7. The pairs are classified by the categories of COGs that are contained. As a COG has more than one COG category and a pair contains COGs belonging to different COG categories, a pair can appear in more than one category. Alphabets in the y-axis represent COG categories as follows. A: RNA processing and modification, C: Energy production and conversion, D: Cell cycle control, cell division, chromosome partitioning, E: Amino acid transport and metabolism, F: Nucleotide transport and metabolism, G: Carbohydrate transport and metabolism, H: Coenzyme transport and metabolism, I: Lipid transport and metabolism, J: Translation, ribosomal structure, and biogenesis, K: Transcription, L: Replication, recombination, and repair, M: Cell wall/membrane/envelope/ biogenesis, N: Cell motility, O: Posttranslational modification, protein turnover, and chaperones, P: Inorganic ion transport and metabolism, Q: Secondary metabolites biogenesis, transport, and metabolism, R: General function prediction only, S: Function unknown, T: Signal transduction mechanism, U: Intercellular trafficking, secretion, and vesicular transport, V: Defense mechanisms, W: Extracellular structure, X: Mobilome, prophages, and transposons. The plots of the other TP thresholds are shown in Supplementary Figures S13 (TPR=0.6) and S14 (TPR=0.5)

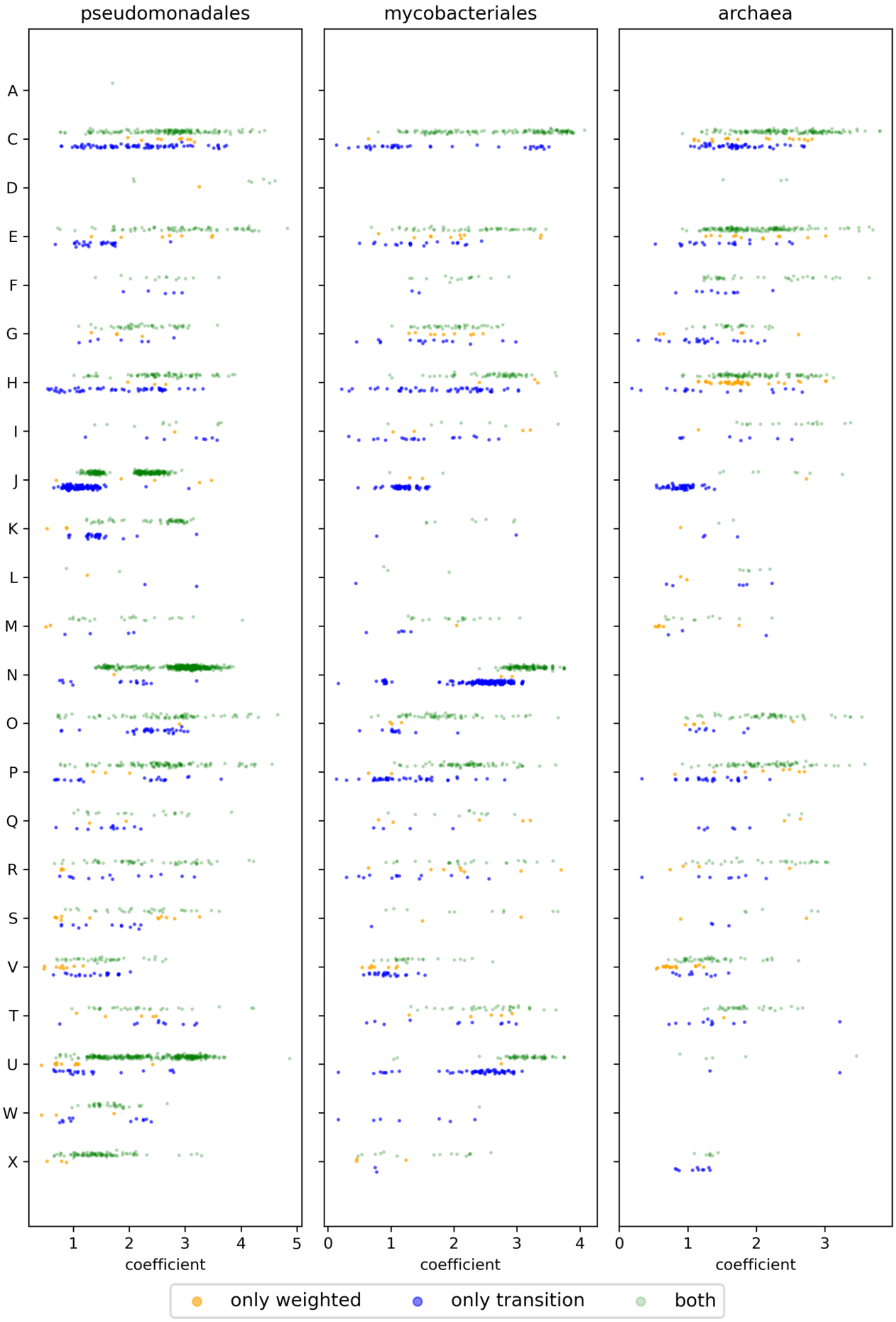

pseudomonadales
mycobacteriales
archaea
A
C
D
E
F
G
H
I
J
K
L
M
N
O
P
Q
R
S
V
T
U
W
X
1
2
3
4
5
0
1
2
3
4
0
1
2
3
coefficient
coefficient
coefficient
only weighted
only transition
both

**Supplementary Figure S7.** The scatter plot of the coefficient (x-axis) and the differences between $t_1$ and $t_2$ in co-evolved pairs in the Pseudomonadales dataset at a True Positive Rate = 0.7. The details are the same as Fig. 3B. The COG categories represented by alphabets are described in the legend of Supplementary Figure S6.

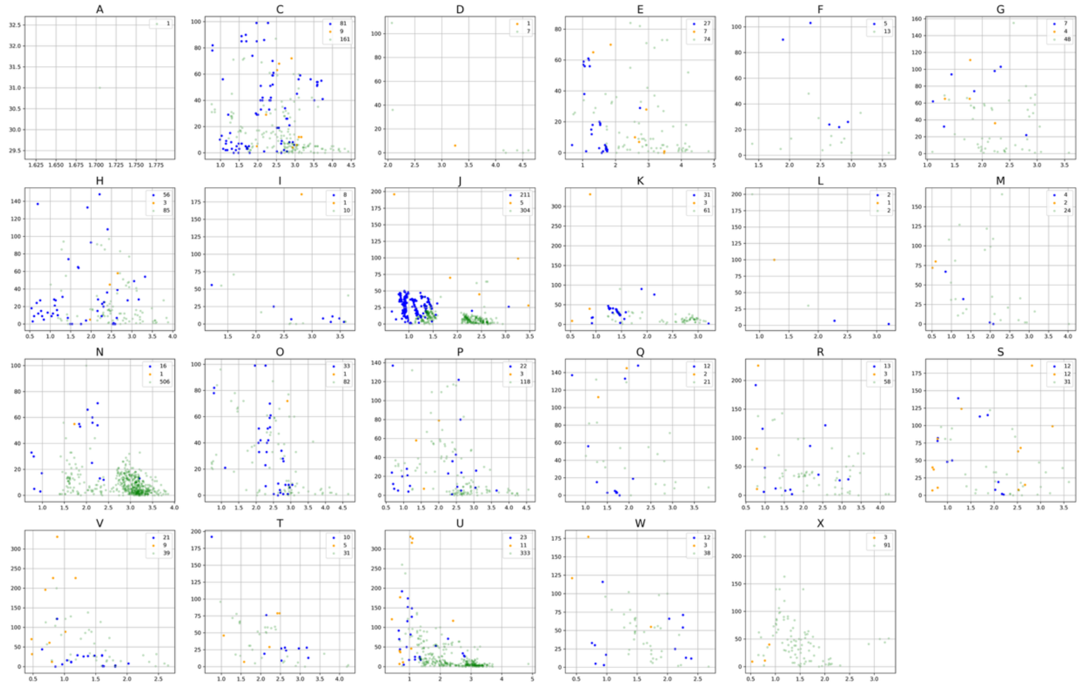

**Supplementary Figure S8.** The scatter plot of the coefficient (x-axis) and the differences between $t_1$ and $t_2$ in co-evolved pairs in the Mycobacteriales dataset at a True Positive Rate = 0.7. The details are the same as Fig. 3B. The COG categories represented by alphabets are described in the legend of Supplementary Figure S6.

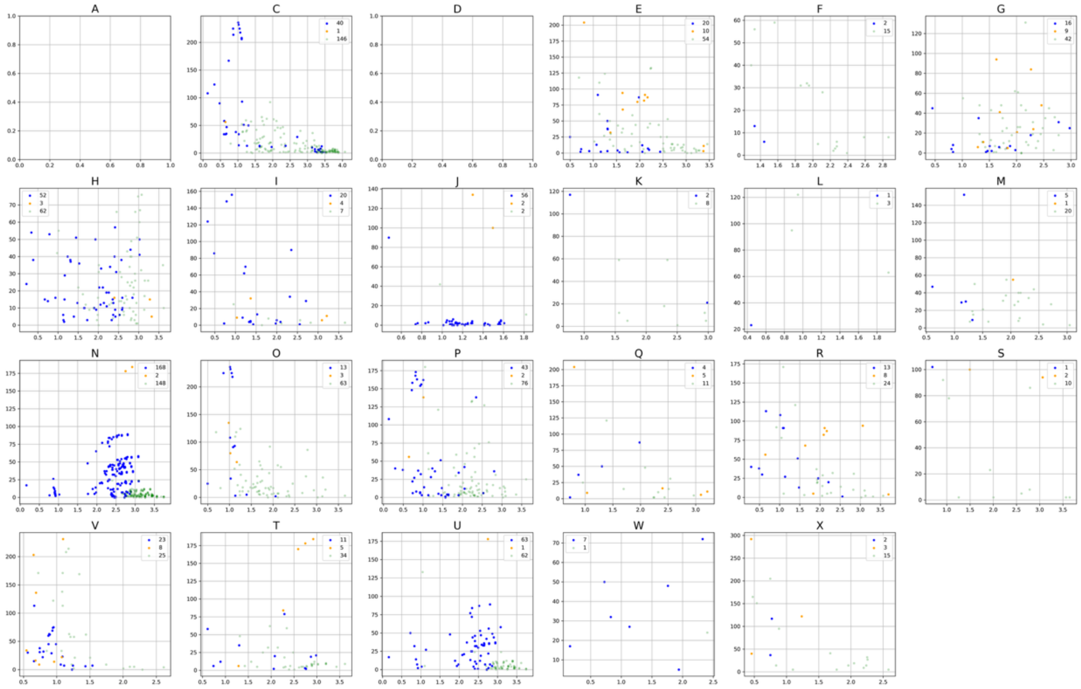

**Supplementary Figure S9.** The scatter plot of the coefficient (x-axis) and the differences between $t_1$ and $t_2$ in co-evolved pairs in the Archaea dataset at a True Positive Rate = 0.7. The details are the same as Fig. 3B. The COG categories represented by alphabets are described in the legend of Supplementary Figure S6.

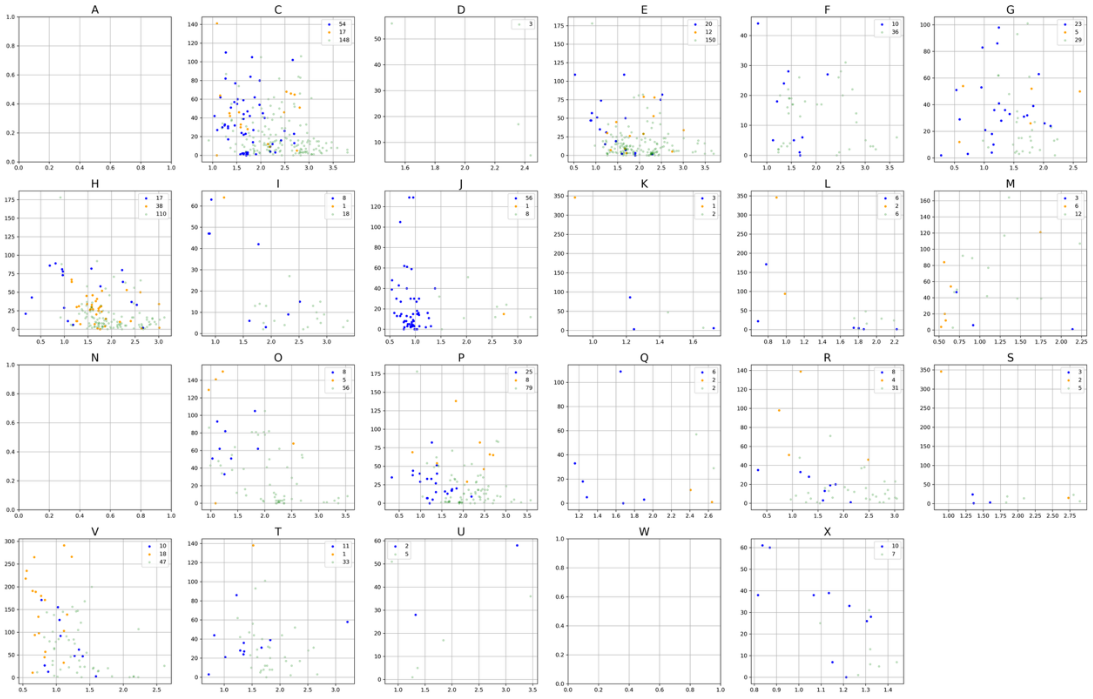

**Supplementary Figure S10.** A network of co-evolved COG pairs detected by phylogenetic profiling in the Mycobacteriales dataset at a True Positive Rate = 0.7. The COGs belonging to category N (cell motility) are connected by the edges when they are detected as co-evolved pairs. The pairs only detected by transition methods are colored in blue and otherwise in green. The fourteen COGs at the bottom only have blue edges, indicating that they were not in the pairs detected by weighted methods.

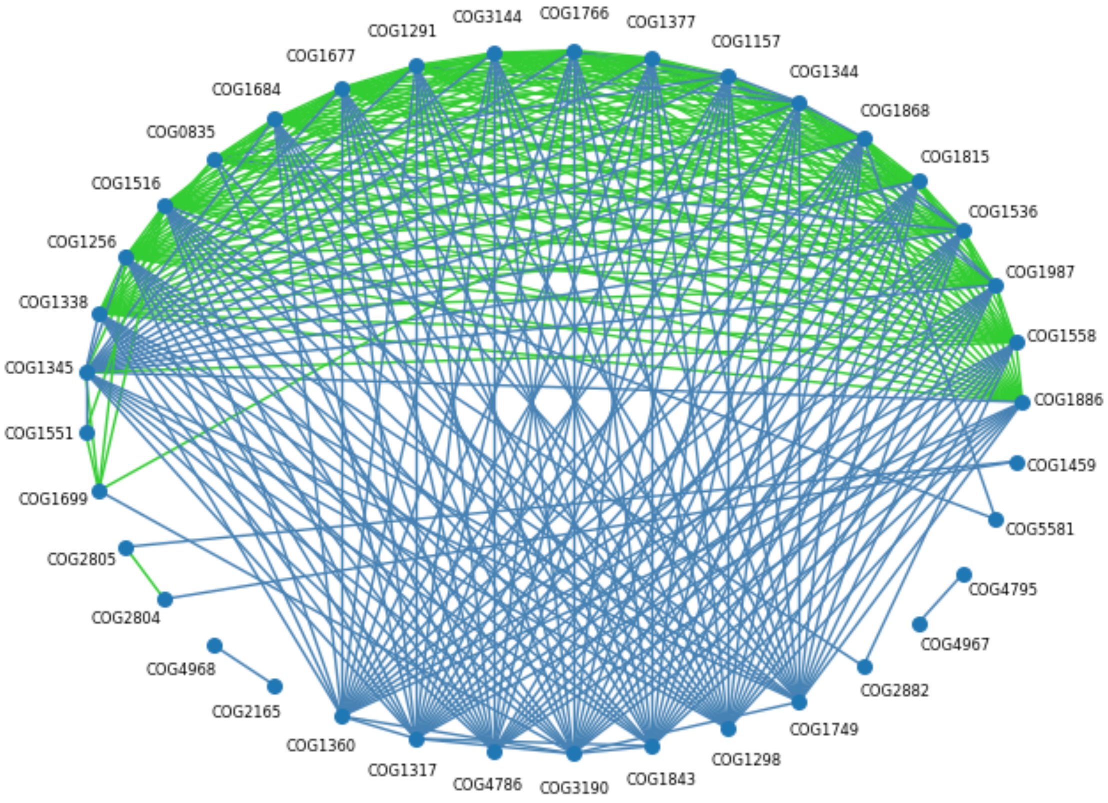

**Supplementary Figure S11.** The EvolCCM coefficient distributions of only detected by weighted methods (orange) and transition methods (blue) at various True Positive Rates (TPRs). The distribution of mycobacteriales at a TPR = 0.9 is not shown as there are a few pairs for kernel density plot.

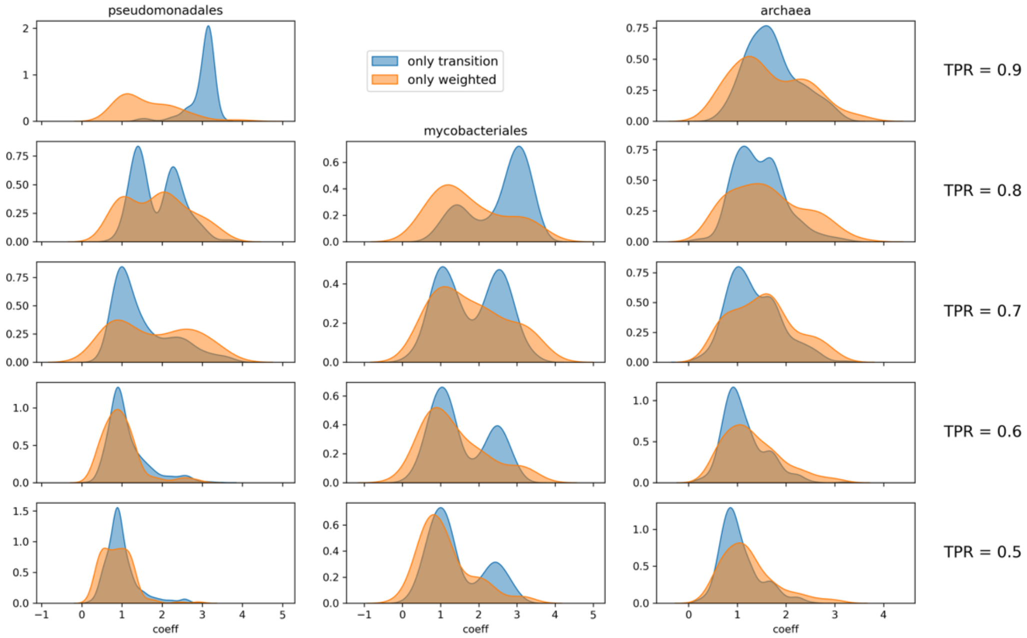

**Supplementary Figure S12.** The scatter plot of the coefficient (x-axis) and the differences between $t_1$ and $t_2$ in co-evolved pairs of COGs belonging to category N (y-axis) in the Pseudomonadales datasets at a True Positive Rate (TPR) = 0.9. $t_1$ and $t_2$ are the number of presence/absence state changes in each COG of the pairs across the phylogenetic tree. Blue dots indicate the co-evolved pairs only detected by transition methods, while green ones indicate pairs detected by both transition and weighted methods.

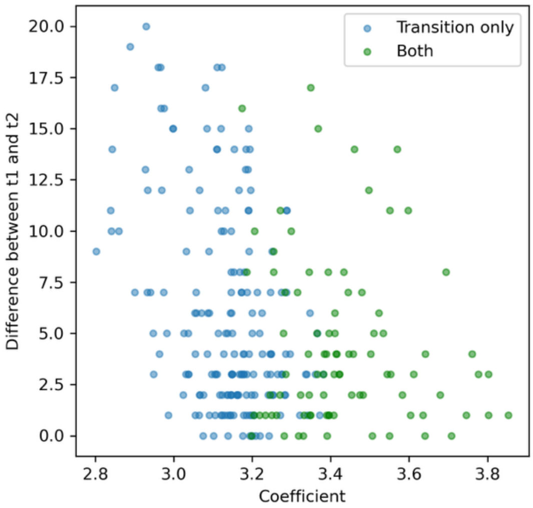

**Supplementary Figure S13.** The distribution of coefficients of the detected positive pairs only detected by transition methods (green), weighted methods (orange), and both (blue) at a True positive Rate (TPR) = 0.6. Details are the same as Supple FigS13_coeff_category06mentary Figure S6.

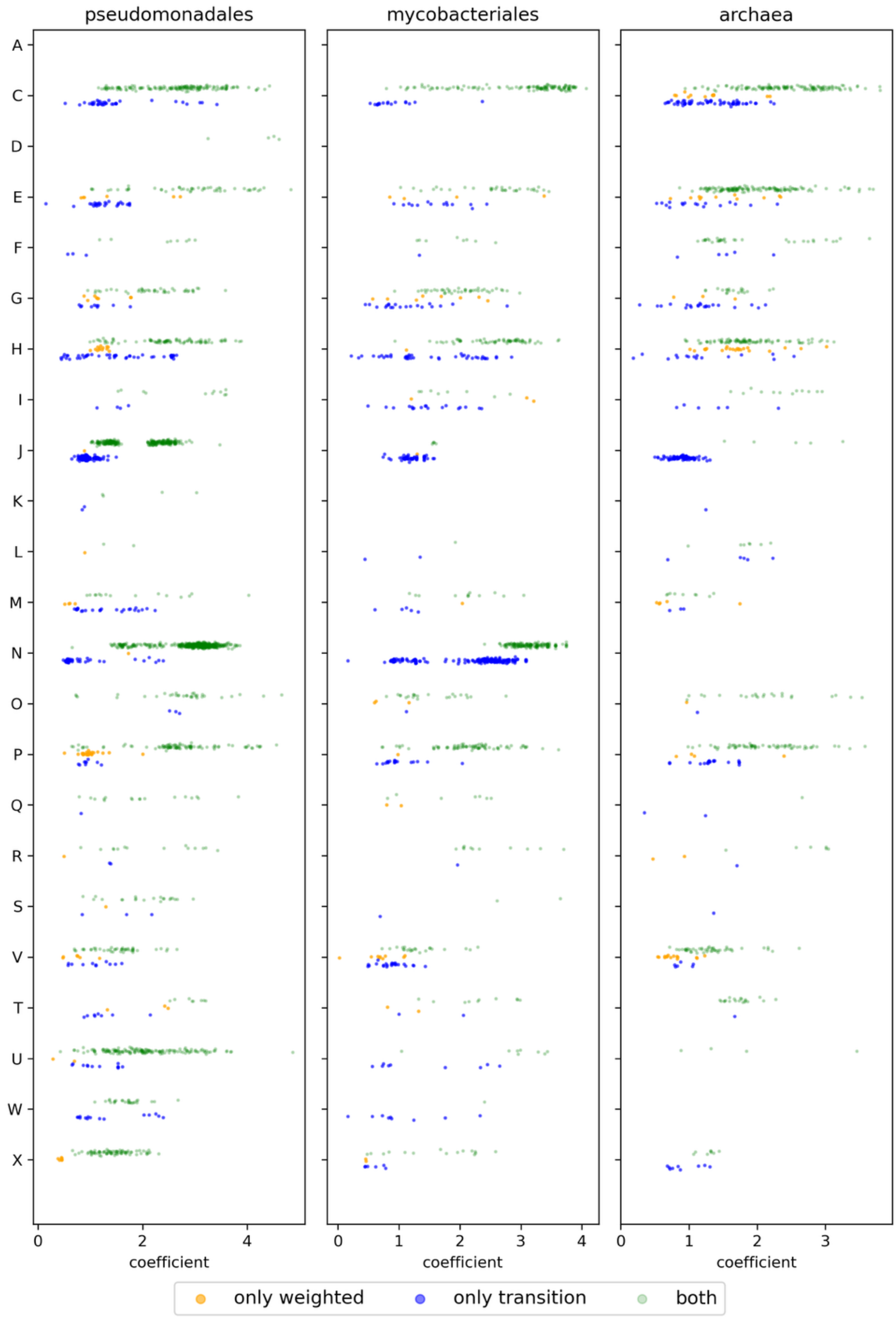

**Supplementary Figure S14.** The distribution of coefficients of the detected positive pairs only detected by transition methods (green), weighted methods (orange), and both (blue) at a True positive Rate (TPR) = 0.5. Details are the same as Supple FigS13_coeff_category06mentary Figure S6.

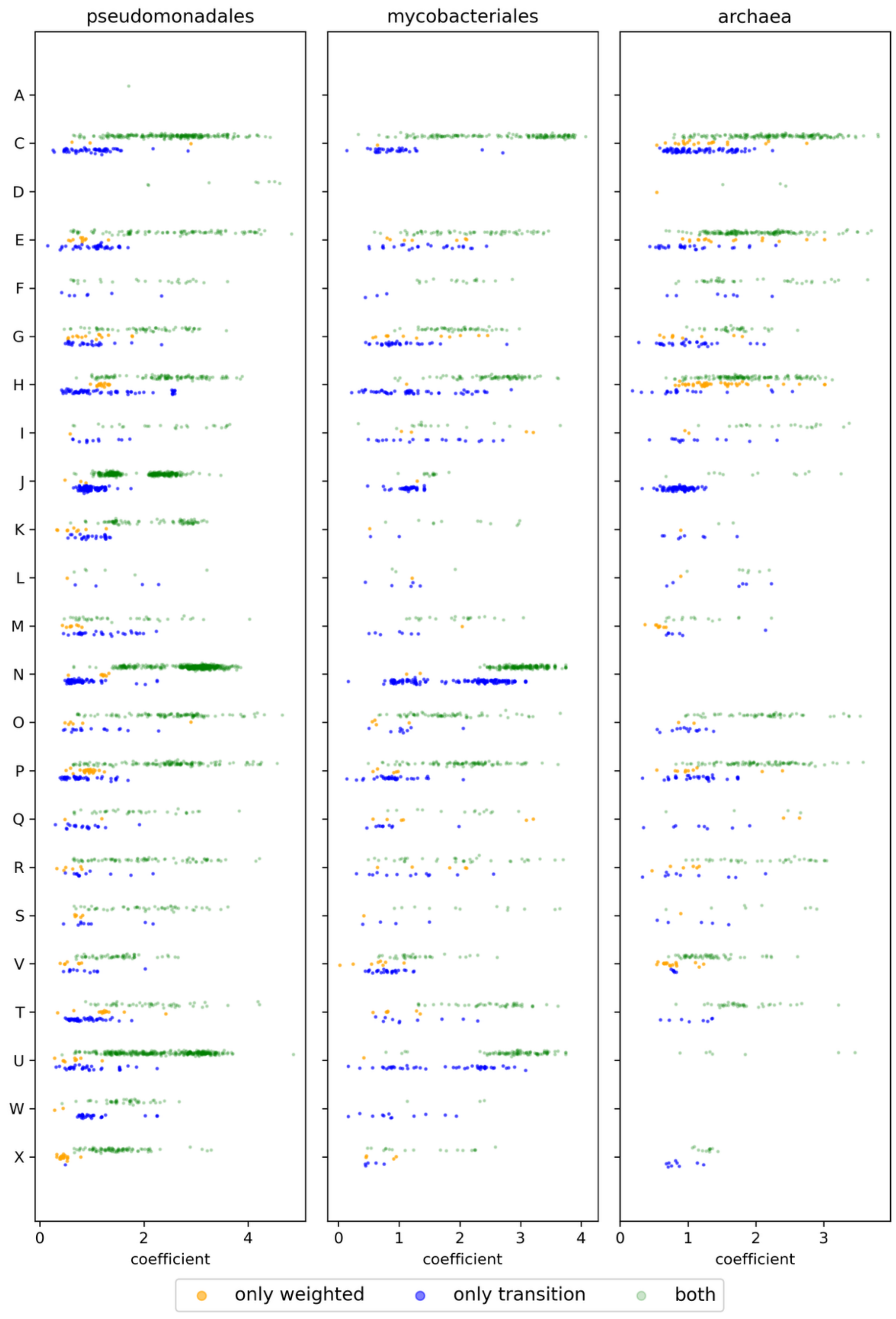

**Supplementary Figure S15.** Comparison of the number of transitions from/to intermediate state between the pairs only detected by transition methods (blue) and weighted methods (orange) at various True Positive Rates (TPR).

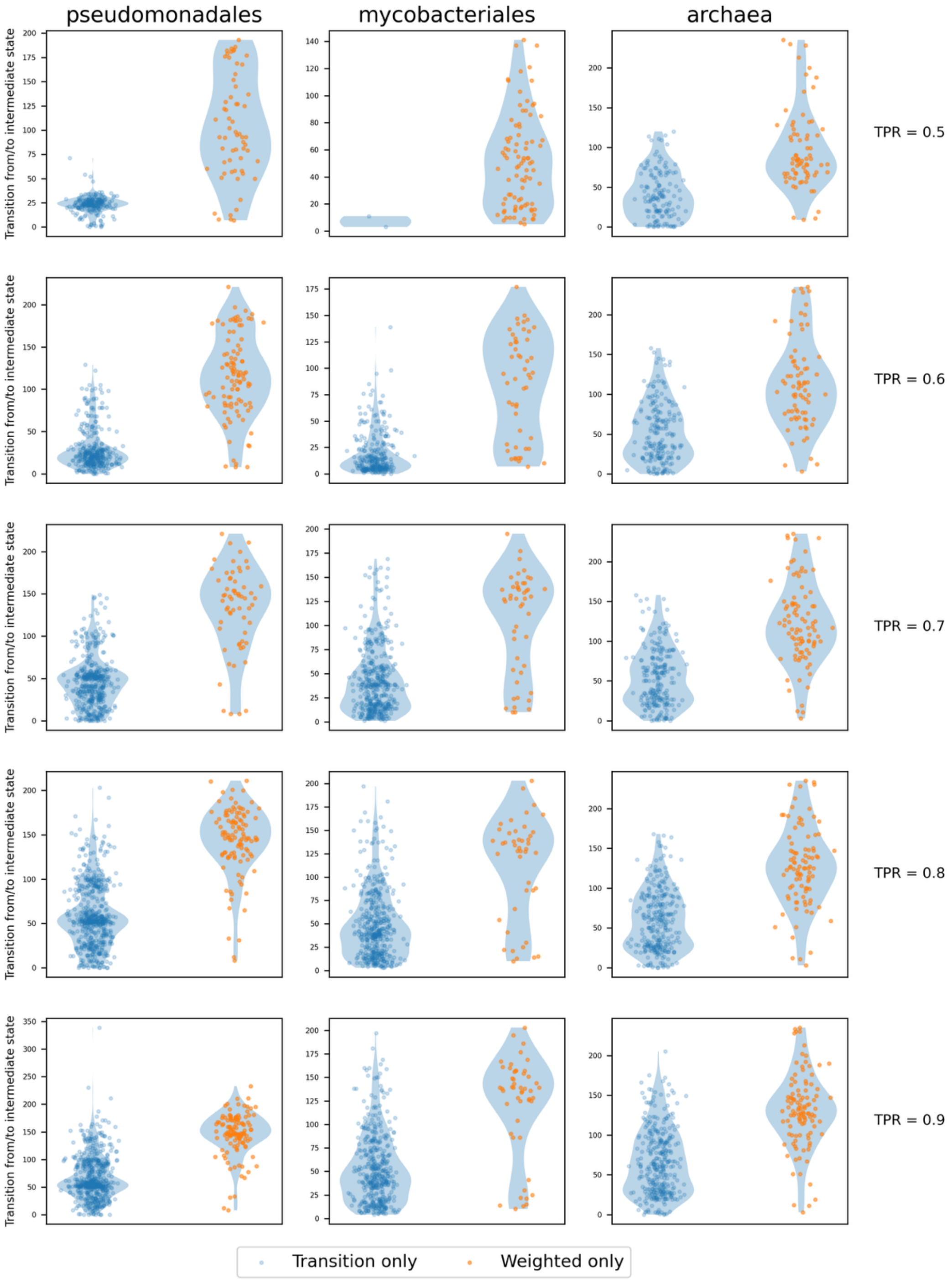

**Supplementary Figure S16.** Characteristic of the weighted methods. Comparison of the numbers of changes from/to the intermediate state of the co-evolved gene pairs only detected by the transition method (blue) and weighted methods (orange) at various True Positive Rates (TPRs). The number of changes was counted from the ancestral state reconstruction result of the co-evolved gene pairs. The numbers of the changes from/to the intermediate state are indicated on the x-axis, and the biases to the changes to one of two intermediate states are indicated on the y-axis. The transition bias was calculated by dividing the larger number of changes from/to the intermediate state of pair genes by their sum.

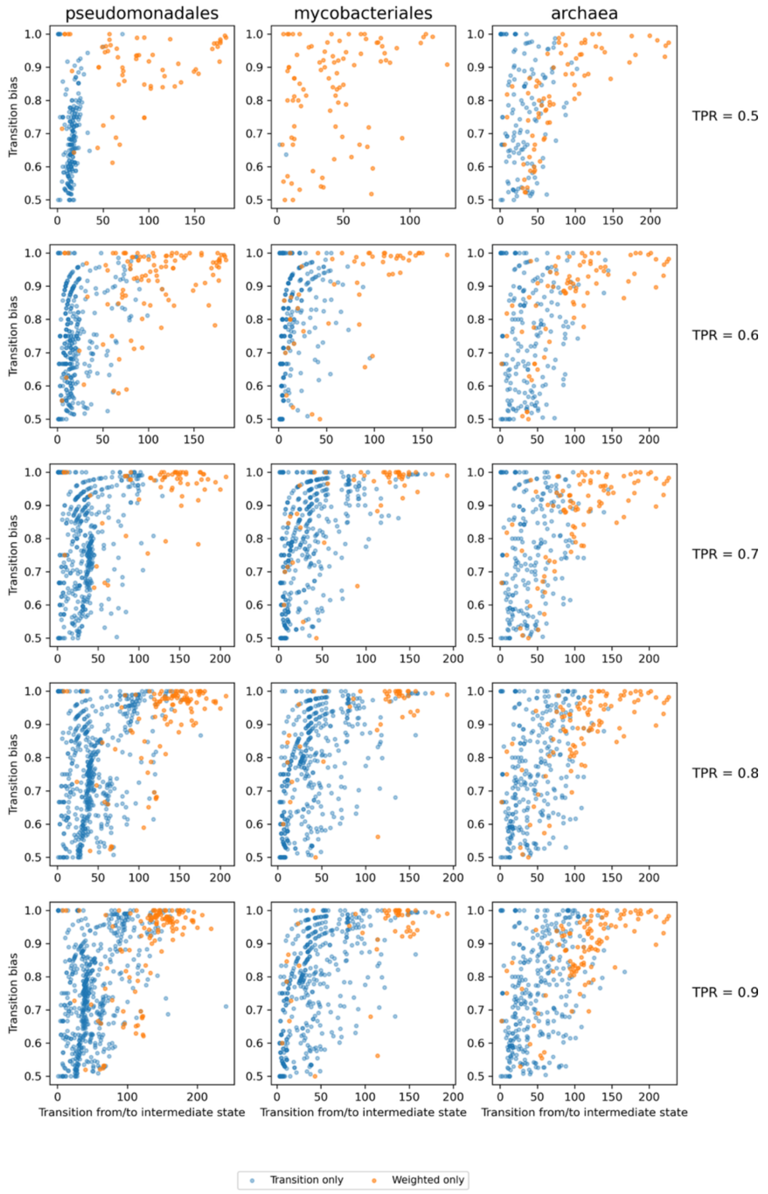

**Supplementary Figure S17.** Precision-Recall (PR) curves of phylogenetic profiling method proposed in this study and those of EvoWeaver. Three prokaryotic datasets were analyzed. Areas under the PR curve (PRAUCs) are shown in parentheses.

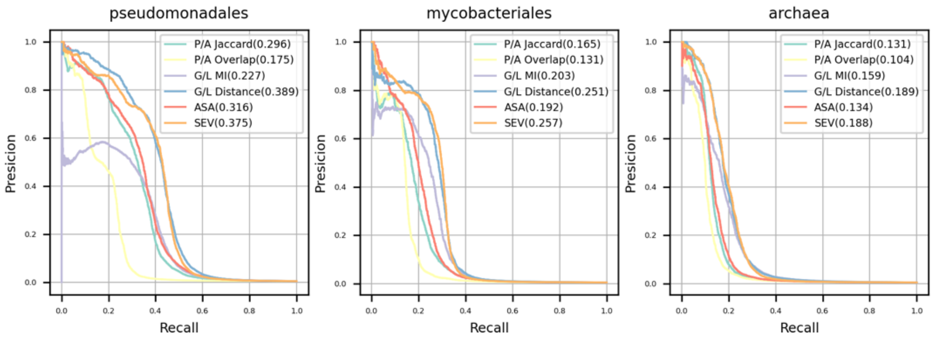


**Supplementary Figure S18.** The number of positive pairs detected by phylogenetic profiling method proposed in this study and those of EvoWeaver. Three prokaryotic datasets were analyzed as well as in Supplementary Figure S17, and the results under the various True Positive Rates are shown.

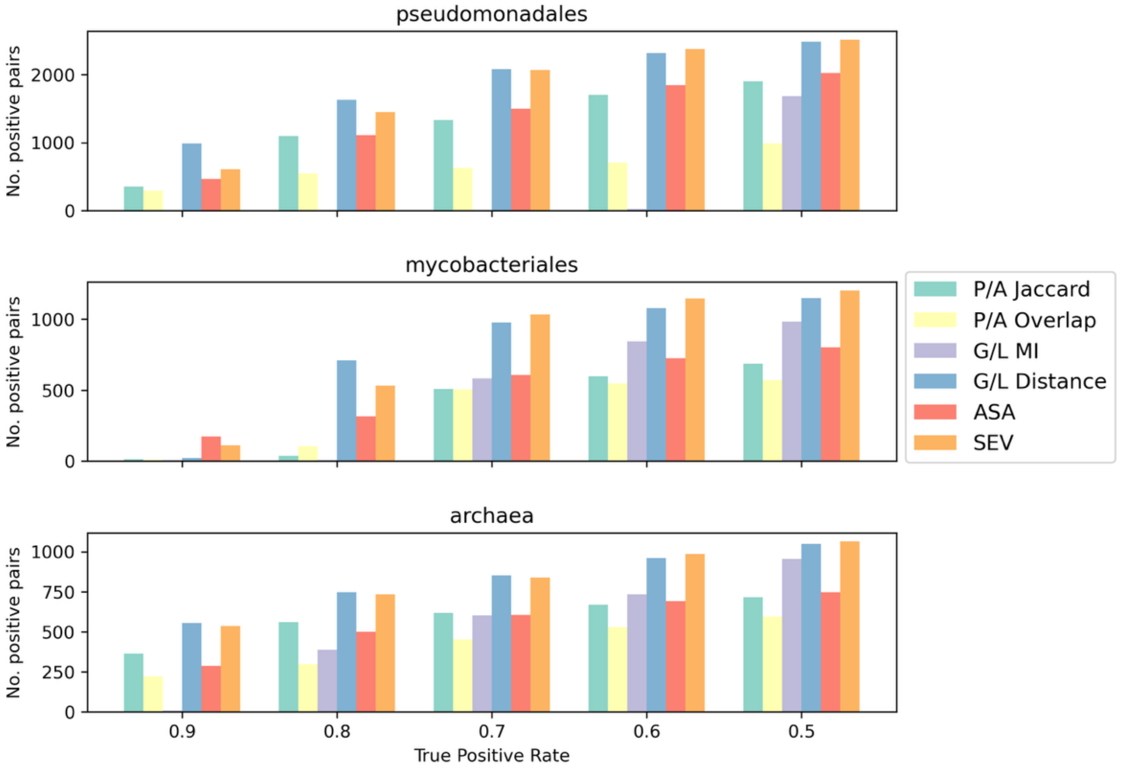

**Supplementary Figure S19.** Comparison of SEV and G/L Distance. Left, the numbers of positive detected pairs at various True Positive Rates are shown in heatmaps. Right, same as the left panel, but the ratios to the number of pairs by labelled methods to those by either of the methods are shown.

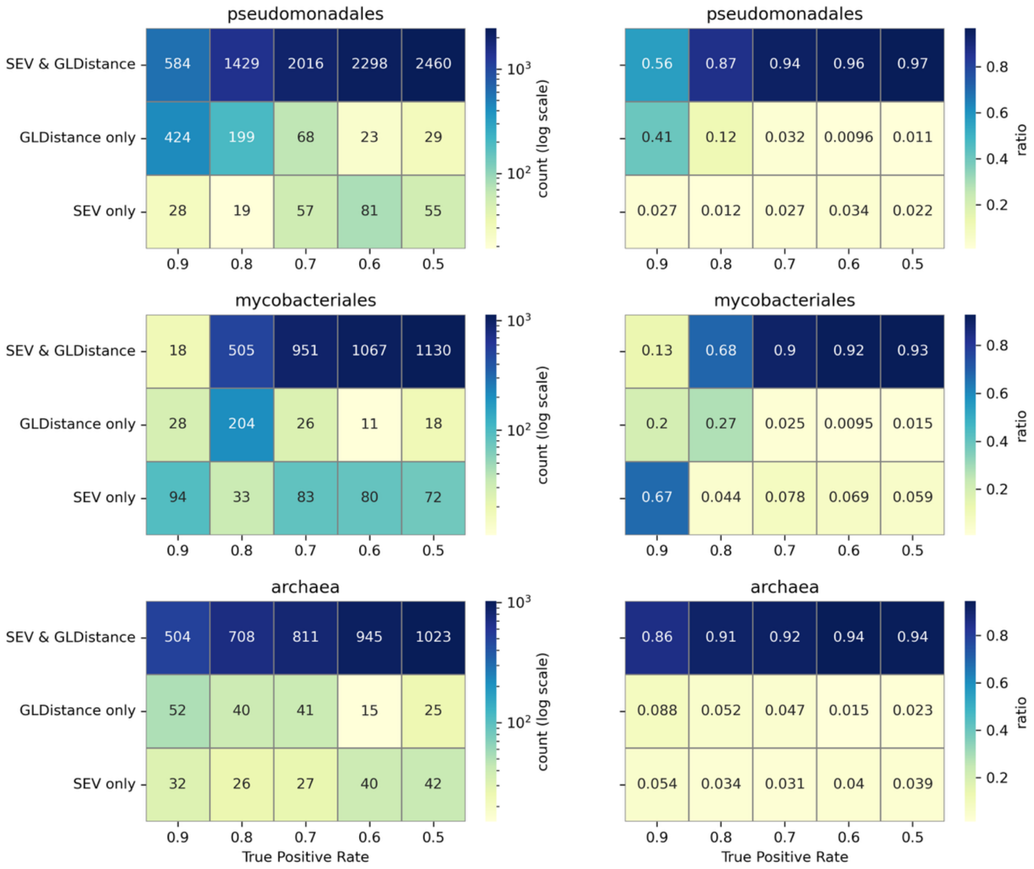
