## Supplementary Tables S1-7 for "CORGIAS: identifying correlated gene pairs by considering evolutionary history in a large-scale prokaryotic genome dataset"

**Supplementary Table S1.** PRAUC of ASA and SEV using various methods for ancestral state reconstruction. "MPPA" is the maximum likelihood method, and the other are the maximum parsimony methods. The best AUC in the dataset and the phylogenetic profiling methods (ASA or SEV) are shown in bold.

| Phylogenetic profiling | Dataset | Ancestral state reconstruction | PRACU |
|---|---|---|---|
| ASA | Pseudomonadasles | MPPA | **0.316** |
| | | DOWNPASS | 0.257 |
| | | ACTRAN | 0.296 |
| | | DELTRAN | 0.290 |
| | Mycobacteriales | MPPA | **0.192** |
| | | DOWNPASS | 0.098 |
| | | ACTRAN | 0.158 |
| | | DELTRAN | 0.146 |
| | Archaea | MPPA | **0.134** |
| | | DOWNPASS | 0.132 |
| | | ACTRAN | 0.133 |
| | | DELTRAN | 0.118 |
| SEV | Pseudomonadasles | DOWNPASS | 0.371 |
| | | ACTRAN | **0.375** |
| | | DELTRAN | 0.370 |
| | Mycobacteriales | DOWNPASS | 0.251 |
| | | ACTRAN | **0.257** |
| | | DELTRAN | 0.251 |
| | Archaea | DOWNPASS | 0.187 |
| | | ACTRAN | **0.188** |
| | | DELTRAN | 0.185 |

**Supplementary Table S2.** The number of positive pairs detected by each phylogenetic profiling method in the Pseudomonadales dataset with varying true positive rates. The number of positive pairs detected by either weighted and transition methods are also shown. The ratios of the number of pairs detected by each method are shown in parentheses.

| **TPR** | **0.9** | **0.8** | **0.7** | **0.6** | **0.5** |
|---|---|---|---|---|---|
| naive | 1 (0.1%) | 99 (6.4%) | 888 (40.9%) | 1073 (42.2%) | 1186 (43.6%) |
| RLE | 358 (49.4%) | 1039 (66.8%) | 1348 (62.0%) | 1588 (62.4%) | 1670 (61.4%) |
| CWA | 265 (36.6%) | 1040 (66.9%) | 1630 (75.0%) | 1844 (72.5%) | 1983 (73.0%) |
| ASA | 465 (64.2%) | 1111 (71.4%) | 1497 (68.9%) | 1847 (72.7%) | 2026 (74.5%) |
| weighted methods | 478 (66.0%) | 1193 (76.7%) | 1706 (78.5%) | 1980 (77.9%) | 2128 (78.3%) |
| cotransitions | **619 (85.5%)** | 1294 (83.2%) | 1966 (90.5%) | 2378 (93.5%) | **2567 (94.4%)** |
| SEV | 609 (84.1%) | **1448 (93.1%)** | **2073 (95.4%)** | **2379 (93.6%)** | 2515 (92.5%) |
| transition methods | 666 (92.0%) | 1457 (93.7%) | 2111 (97.1%) | 2432 (95.7%) | 2595 (95.5%) |
| Total | 724 | 1555 | 2173 | 2542 | 2718 |

**Supplementary Table S3.** The number of positive pairs detected by each phylogenetic profiling method in the Mycobacteriales dataset with varying true positive rates. The details are the same as Supplementary Table S3

| TPR | 0.9 | 0.8 | 0.7 | 0.6 | 0.5 |
|---|---|---|---|---|---|
| naive | 85 (42.7%) | 221 (34.1%) | 320 (28.9%) | 439 (35.6%) | 509 (39.5%) |
| RLE | 127 (63.8%) | 251 (38.7%) | 303 (27.3%) | 334 (27.1%) | 342 (26.5%) |
| CWA | 123 (61.8%) | 298 (45.9%) | 365 (32.9%) | 435 (35.3%) | 475 (36.8%) |
| ASA | **174 (87.4%)** | 316 (48.7%) | 607 (54.8%) | 676 (58.9%) | 802 (62.1%) |
| weighted methods | 195 (98.0%) | 421 (63.5%) | 671 (60.1%) | 784 (63.6%) | 856 (66.4%) |
| cotransitions | 107 (53.8%) | 514 (79.2%) | 988 (89.2%) | 1124 (91.2%) | 1192 (92.4%) |
| SEV | 112 (56.3%) | **533 (82.1%)** | **1034 (93.3%)** | **1147 (92.4%)** | **1202 (93.1%)** |
| transition methods | 117 (58.8%) | 579 (89.2%) | 1056 (95.3%) | 1184 (96.0%) | 1237 (95.9%) |
| Total | 199 | 649 | 1108 | 1233 | 1290 |

**Supplementary Table S4.** The number of positive pairs detected by each phylogenetic profiling method in the Archaea dataset with varying true positive rates. The details are the same as Supplementary Table S3

| **TPR** | **0.9** | **0.8** | **0.7** | **0.6** | **0.5** |
|---|---|---|---|---|---|
| naive | 23 (3.7%) | 272 (32.5%) | 365 (38.1%) | 418 (37.8%) | 485 (40.3%) |
| RLE | 326 (53.6%) | 372 (44.4%) | 396 (41.3%) | 426 (38.7%) | 447 (37.1%) |
| CWA | 363 (59.7%) | 469 (56.0%) | 527 (55.0%) | 590 (53.6%) | 667 (55.4%) |
| ASA | 287 (47.2%) | 501 (59.9%) | 605 (63.1%) | 691 (62.8%) | 748 (62.1%) |
| weighted methods | 456 (75.0%) | 633 (75.6%) | 724 (75.5%) | 807 (73.3%) | 865 (71.8%) |
| cotransitions | 515 (84.7%) | 708 (84.6%) | 805 (83.9%) | 889 (80.7%) | 1019 (84.6%) |
| SEV | **536 (88.2%)** | **734 (87.7%)** | **838 (87.4%)** | **985 (89.5%)** | **1065 (88.5%)** |
| transition methods | 556 (91.4%) | 763 (91.2%) | 867 (90.4%) | 1005 (91.2%) | 1095 (90.9%) |
| Total | 199 | 649 | 1108 | 1233 | 1290 |

**Supplementary Table S5.** The number of positive pairs only identified by each phylogenetic profiling method in the Pseudomonadales dataset with varying true positive rates. The number of positive pairs detected by either weighted methods but not the transition methods, and vice versa, are also shown. The ratios of the number of pairs detected by each method are shown in parentheses.

| **TPR** | **0.9** | **0.8** | **0.7** | **0.6** | **0.5** |
|---|---|---|---|---|---|
| naive | 0 (0%) | 3 (0.2%) | 5 (0.2%) | 7 (0.3%) | 5 (0.2%) |
| RLE | 5 (0.7%) | 7 (0.5%) | 7 (0.3%) | 11 (0.4%) | 7 (0.3%) |
| CWA | 0 (0%) | 5 (0.3%) | 11 (0.5%) | 21 (0.8%) | 29 (1.0%) |
| ASA | 22 (3.0%) | 24 (1.5%) | 4 (0.2%) | 24 (0.9%) | 24 (0.9%) |
| weighted methods | 58 (8.0%) | 98 (6.3%) | 62 (2.9%) | 110 (4.3%) | 123 (4.5%) |
| cotransitions | 52 (7.2%) | 5 (0.3%) | 19 (0.9%) | 30 (1.2%) | 49 (1.8%) |
| SEV | 42 (5.8%) | 150 (9.6%) | 144 (6.6%) | 52 (2.0%) | 26 (1.0%) |
| transition methods | 246 (34%) | 362 (23.2%) | 467 (21.4%) | 562 (22.1%) | 590 (21.7%) |

**Supplementary Table S6.** The number of positive pairs only identified by each phylogenetic profiling method in the Mycobacteriales dataset with varying true positive rates. The details are the same as Supplementary Table S3.

| **TPR** | **0.9** | **0.8** | **0.7** | **0.6** | **0.5** |
|---|---|---|---|---|---|
| naive | 16 (8.0%) | 23 (3.5%) | 10 (0.9%) | 6 (0.5%) | 5 (0.4%) |
| RLE | 3 (1.5%) | 9 (1.4%) | 9 (0.8%) | 9 (0.7%) | 8 (0.6%) |
| CWA | 1 (0.5%) | 8 (1.2%) | 5 (0.5%) | 5 (0.4%) | 10 (0.8%) |
| ASA | 18 (9.0%) | 5 (0.8%) | 12 (1.0%) | 7 (0.6%) | 12 (1.0%) |
| weighted methods | 82 (41.2%) | 70 (1.1%) | 52 (4.7%) | 49 (4.0%) | 53 (4.1%) |
| cotransitions | 0 (0%) | 32 (5.0%) | 11 (1.0%) | 24 (2.0%) | 25 (1.9%) |
| SEV | 3 (1.5%) | 59 (9.0%) | 61 (5.5%) | 58 (4.7%) | 43 (3.3%) |
| transition methods | 4 (2.0%) | 237 (36.5%) | 437 (39.4%) | 449 (36.4%) | 434 (33.6%) |

**Supplementary Table S7.** The number of positive pairs only identified by each phylogenetic profiling method in the Archaea dataset with varying true positive rates. The details are the same as Supplementary Table S3.

| TPR | 0.9 | 0.8 | 0.7 | 0.6 | 0.5 |
|---|---|---|---|---|---|
| naive | 5 (0.8%) | 13 (1.5%) | 17 (1.8%) | 12 (1.1%) | 17 (1.4%) |
| RLE | 12 (2.0%) | 13 (1.5%) | 13 (1.4%) | 12 (1.1%) | 10 (0.8%) |
| CWA | 16 (2.6%) | 19 (2.3%) | 10 (1.0%) | 16 (1.5%) | 16 (1.3%) |
| ASA | 3 (0.5%) | 15 (1.8%) | 24 (2.5%) | 31 (2.8%) | 26 (2.1%) |
| weighted methods | 52 (8.6%) | 74 (8.8%) | 92 (9.6%) | 96 (4.0%) | 109 (9.1%) |
| cotransitions | 8 (1.3%) | 14 (1.7%) | 15 (1.6%) | 6 (0.5%) | 16 (1.3%) |
| SEV | 34 (5.6%) | 48 (5.7%) | 54 (5.6%) | 103 (9.4%) | 74 (6.1%) |
| transition methods | 152 (25.0%) | 204 (24.4%) | 235 (24.5%) | 294 (36.4%) | 339 (28.2%) |
