## Supplementary Table 11 for "CORGIAS: identifying correlated gene pairs by considering evolutionary history in a large-scale prokaryotic genome dataset"

**Supplementary Table S11.** Runtime regression of the phylogenetic profiling methods used in this study. Runtimes (t) were regressed on the following: log(t) = $\alpha_1$log(number of species) + $\alpha_2$log(number of genes) + $\beta$. The predicted runtimes of the latest GTDB (Release 220, 107,000 species and 6,000 OGs) and OrthoDB (v12, 28,000 species, and 61,000 OGs) are also shown. The numbers of OGs were obtained from previous studies (Tremblay *et al.* 2021, Dembech *et al.* 2023).

| | $\alpha_1$ | $\alpha_2$ | $\beta$ | $R^2$ | OrthoDB | GTDB |
|---|---|---|---|---|---|---|
| naïve | 1.52 | 2.23 | -25.6 | 0.990 | 31 d | 54 h |
| naïve w/ GPU | 0.219 | 1.52 | -12.2 | 0.979 | 5.8 m | 10 m |
| RLE | 0.205 | 2.02 | -12.0 | 0.999 | 19 h | 51 h |
| CWA | 0.905 | 2.00 | -15.4 | 0.997 | 63 d | 22 d |
| ASA | 1.09 | 1.85 | -13.2 | 0.999 | 2.9 y | 195 d |
| cotransitions | 0.847 | 1.79 | -17.6 | 0.982 | 10.7 h | 3.8 h |
| cotransitions w/ GPU | 0.336 | 1.27 | -10.7 | 0.951 | 8.3 m | 8.4 m |
| SEV | 0.884 | 1.851 | -17.7 | 0.998 | 27 h | 9.0 h |
| SEV w/ GPU | 0.521 | 1.61 | -14.5 | 0.954 | 48 m | 38 m |
| G/L Distance | 1.091 | 2.01 | -17.2 | 0.992 | 52 d | 11d |
